# Interrogation of cancer gene dependencies reveals novel paralog interactions of autosome and sex chromosome encoded genes

**DOI:** 10.1101/2021.05.21.445116

**Authors:** Anna Köferle, Andreas Schlattl, Alexandra Hörmann, Fiona Spreitzer, Alexandra Popa, Venu Thatikonda, Teresa Puchner, Sarah Oberndorfer, Corinna Wieshofer, Maja Corcokovic, Christoph Reiser, Simon Wöhrle, Johannes Popow, Mark Pearson, Barbara Mair, Ralph A. Neumüller

## Abstract

Genetic networks are characterized by extensive buffering. During tumour evolution, disruption of these functional redundancies can create *de novo* vulnerabilities that are specific to cancer cells. In this regard, paralog genes are of particular interest, as the loss of one paralog gene can render tumour cells dependent on a remaining paralog. To systematically identify cancer-relevant paralog dependencies, we searched for candidate dependencies using CRISPR screens and publicly available loss-of-function datasets. Our analysis revealed >2,000 potential candidate dependencies, several of which were subsequently experimentally validated. We provide evidence that *DNAJC15-DNAJC19, FAM50A-FAM50B* and *RPP25-RPP25L* are novel cancer relevant paralog dependencies. Importantly, our analysis also revealed unexpected redundancies between sex chromosome genes. We show that chrX- and chrY- encoded paralogs, as exemplified by *ZFX-ZFY, DDX3X-DDX3Y* and *EIF1AX-EIF1AY*, are functionally linked so that tumour cell lines from male patients with Y-chromosome loss become exquisitely dependent on the chrX-encoded gene. We therefore propose genetic redundancies between chrX- and chrY- encoded paralogs as a general therapeutic strategy for human tumours that have lost the Y-chromosome.

## Introduction

Paralog genes that fulfill similar functions provides a degree of robustness of gene regulatory networks to deleterious events^1–3^. These paralog genes arise as a result of gene duplications and subsequent divergent evolution^4,5^. Paralog redundancies are of interest to cancer biology, as tumour-specific processes like hypermethylation, mutations or copy number alterations can inactivate genes and thereby reduce the extent of genetic buffering. Genes whose loss is buffered in non-neoplastic cells by a paralog can thus become dependencies in tumours where the redundant paralog is absent. Examples for such paralog dependencies in cancer cells have been identified and validated before, and include *ENO1-ENO2*^6^, *SMARCA2-SMARCA4*^7–9^, *ARID1A-ARID1B*^10^ or *STAG1-STAG2*^11,12^.

In all validated cases, the tumour-specific loss of one paralog gene creates a specific dependency on a remaining paralog. Accordingly, therapeutic inhibition of the remaining paralog gene is assumed to be safe, because non-tumour cells still retain the genetic buffer to tolerate the inhibition without systemic side effects. Another advantage of tumour-specific vulnerabilities created by loss of a paralogous gene is the availability of a tractable biomarker; i.e. measuring loss of Paralog A in tumours allows to select patients that would respond to inhibition of the synthetic lethal Paralog B. Therefore, paralog dependencies represent a highly attractive concept for cancer drug target identification. However, a systematic understanding of cancer-relevant paralog dependencies is still elusive to date, although CRISPR-based combinatorial screens and bioinformatics discovery pipelines are beginning to shed light on the tumour redundancy map^3,13–20^.

In addition to mutagenic processes such as gene silencing, point mutations or gene amplification, human cancers frequently lose large amounts of their genetic material during the process of tumourigenesis^21^. Deletions can involve one or multiple genes or, as is being appreciated, extend to loss of whole chromosomes, one of the most prevalent being loss of chromosome Y (LOY). Cancer incidence is generally higher in males^22,23^, a fact that has been attributed to the general protective effect of the chrXX status in females that allows buffering of deleterious mutations^24,25^. LOY has been reported in ~93% of esophageal adenocarcinomas^26^, ~12% of male breast cancers^27^ and ~23% of urothelial bladder cancer samples^28^. Mosaic loss of chrY has also been observed outside of the oncology context, where it has been correlated with increased age^29–31^. LOY has been associated with a number of pathophysiological conditions, including clonal hematopoiesis and Alzheimer’s disease^32,33^. Due to the plethora of disease states in which LOY occurs, strategies to eliminate cells involved in pathological conditions, including neoplastic transformation is clearly of general medical interest.

We set out to discover cancer-relevant paralog dependencies by an integrative approach of combining multiple-omics datasets in panels of human cancer cell lines. This analysis reveals 2,040 candidate paralog gene interactions, a subset of which we validate experimentally. Importantly, we uncover a sex-chromosome-specific set of genes that is functionally buffered between the X and Y chromosomes. We demonstrate that targeted depletion of the chrX-encoded gene in LOY tumour cell lines offers an attractive strategy to treat tumours that have lost chrY. In addition, these results provide a generalizable framework of how to eliminate putative pre-pathogenic LOY cells.

## Results

### CRISPR/Cas9 screens identify *CSTF2-CSTF2T* as a paralog dependency

We devised a set of proof-of-principle CRISPR screens to investigate the concept of cancer-specific paralog dependencies. We started by cataloging potential human paralog genes from Ensembl BioMarT, as defined by all genes with at least one other paralog in the same family without any further constraints on homology or family size. Most of the multi-gene families contain two to five paralog genes and the biggest class of genes are protein-coding genes (Figure 1a,b). A compact protein domain-focused CRISPR gRNA library^34^ of ~10,000 gRNAs was generated to permit screening across multiple tumour cell lines. The library genes were manually curated to contain genes that are frequently deleted in human solid tumours, including 460 unique paralog genes from 199 families. Loss-Of-Function (LOF) screens were then carried out across seven cancer cell lines (Hep 3B2.1-7, HuP-T4, MIA PaCa-2, NCI-H1373, NCI-H1993, NCI-H2009, PC-9) with annotated deep deletions (see Methods).

**Figure 1:**
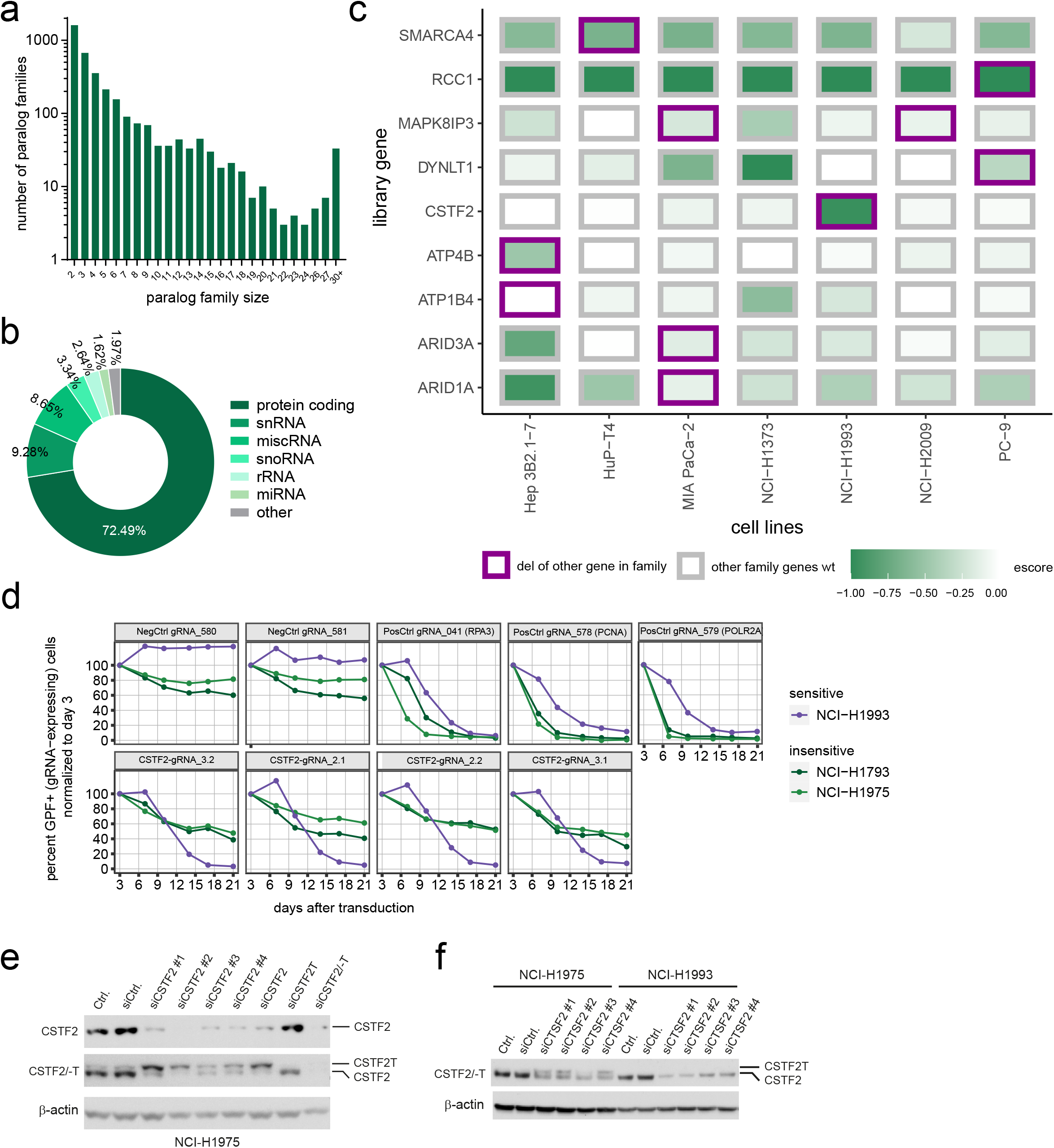
Proof-of-concept paralog dependency CRISPR screens reveal a functional interaction of *CSTF2* and *CSTF2T*. a-b) Distribution of paralog families in study by family size (a) and gene type (b). c) CRISPR screen results in 7 cancer cell lines. Only genes with escore < −0.4 and FDR < 0.1 are displayed. Shades of green indicate effect size (escore), box color indicates whether paralog family contains deleted gene different from listed gene (del) or not (wt). d) CRISPR/Cas9 depletion assay in cell lines resistant (green) and sensitive (purple) to loss of *CSTF2*. gRNAs targeting positive control genes (*RPA3, POLR2A, PCNA*), negative controls (*NegCon03/-07*), and *CSTF2* are indicated. Cells were lentivirally transduced with the gRNA plasmid containing GFP; GFP percentage in transduced cell line pool was measured by flow cytometry at the indicated time points and normalized to day 3 post-transduction. e-f) Western blots for *CSTF2* and *CSTF2T* in lysates from indicated cell lines after siRNA treatment (3d). β-actin was used as loading control.

Paralog genes scoring as significantly depleted in our screens were cross-referenced across the different cell lines to understand whether any of their family members were annotated as deleted (Figure 1c, Supplementary Table 1). Accordingly, we found that *ATP4B* was specifically required in Hep 3B2.1-7 cells that harbor a deletion of *ATP1B2*. Both genes are subunits of potassium-transporting ATPases, but physical or functional interactions have not been described. Furthermore, we observed that NCI-H1993 cells were particularly sensitive to loss of *CSTF2*, likely due to a deletion of its paralog *CSTF2T*. *CSTF2* and *CSTF2T* encode the CstF-64 and CstF-64tau proteins respectively, that have partially overlapping functions in the Cleavage stimulation Factor (CstF) complex, a regulatory component of the mRNA cleavage and polyadenylation machinery^35–37^. We confirmed the sensitivity of *CSTF2T*-negative cells to depletion of *CSTF2* (Figure 1d, Supplementary Figure 1). Mechanistically, we observed that depletion of CSTF2 leads to compensatory induction of *CSTF2T* in *CSTF2T*-proficient cells (Figure 1e), confirming previous reports describing that *CSTF2* and *CSTF2T* can regulate each other’s expression^36–38^. This compensatory upregulation is not observed in *CSTF2T*-deficient cell lines (Figure 2f), providing a hypothesis for the dependency on *CSTF2*. Of note, *CSTF2* and *CSTF2T* are encoded by a single essential gene in yeast, *RNA15 (YGL044C)*^39^, suggesting that cellular viability might depend on the activity of both paralogues in mice and humans. In mice, previous studies suggest that *Cstf2* and *Cstf2t* form a functionally redundant pair of genes with an essential function in certain contexts. Embryonic stem cells lacking *Cstf2* had altered pluripotency and could be differentiated into mesoderm and ectoderm, but not endoderm^38,40^. Autosomally encoded *CSTF2T* is required in pachytene spermatocytes to overcome the lack of expression of X-encoded *CSTF2* due to meiotic sex chromosome inactivation, leading to male sterility^35^. Hence, some but not all functions of Cstf2 can be assumed by Cstf2t.

**Figure 2:**
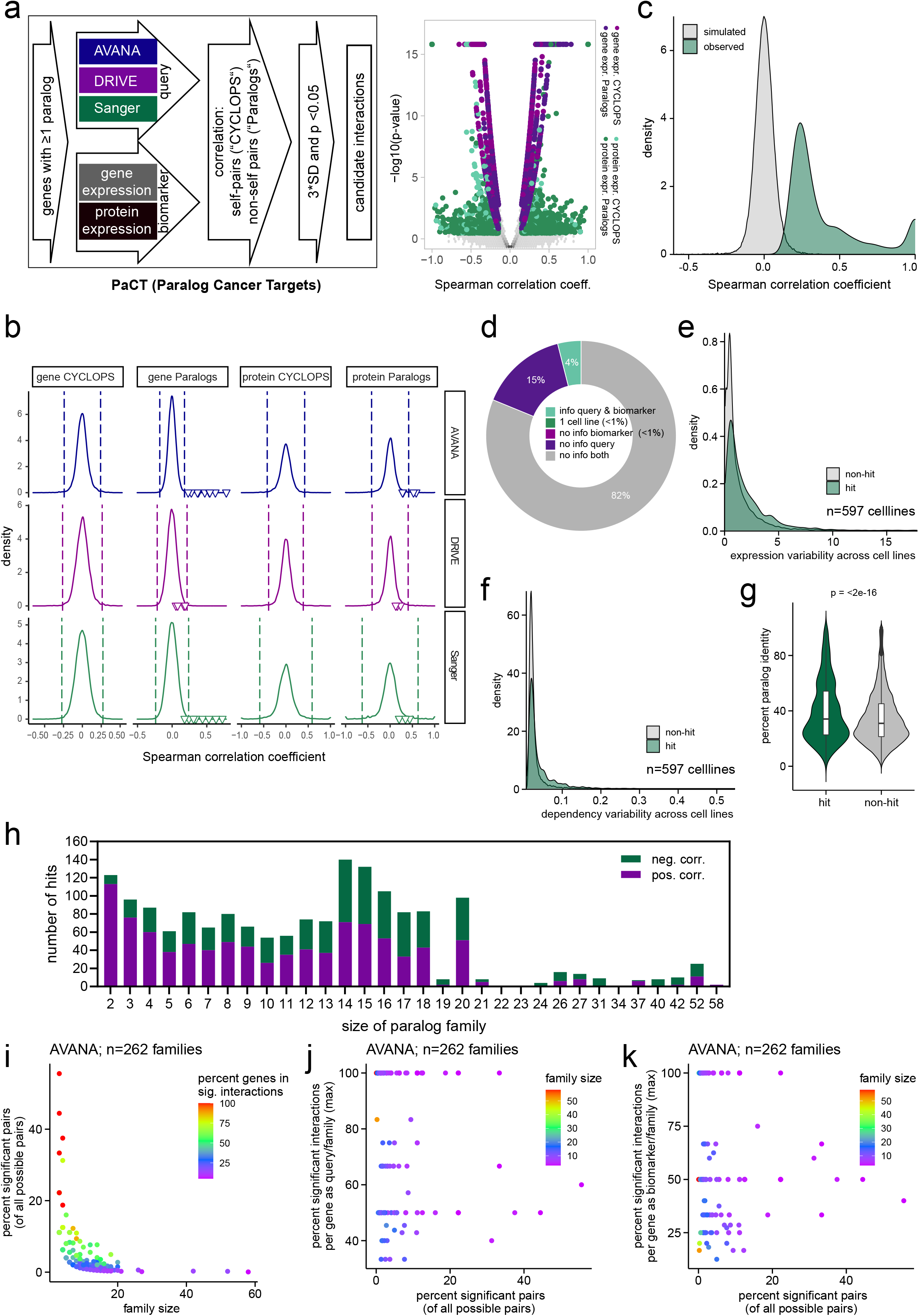
Correlation analysis of public loss-of-function screens yield to identify paralog genetic interactions. a) PaCT analysis workflow and volcano plot of tested pairs by dataset (CYCLOPS, self-interactions; Paralogs, pairwise paralog interactions within same family). See Methods for details. b) Distribution of PaCT correlations (Spearman) by input datasets. Triangles indicate specific hit pairs mentioned in subsequent analyses. Dashed lines mark the 3-standard-deviations cutoff used for hit filtering. c) Distribution of Spearman correlation coefficients of randomly assigned genes to each query compared to the correlation distribution of original PaCT hits. d) Pie chart displaying different categories of query-biomarker pairs across the complete theoretical PaCT exploratory space. Only pairs for which information for both query and biomarker is available can yield hit interactions. See Methods for details. e, f) Expression (e) and depletion score (f) variability distribution of genes involved in hit and non-hit interactions across cell lines. g) Nucleotide sequence similarity difference between hit and non-hit pairs. h) Number of unique hit pairs in paralog families of different size, grouped by type of correlation. i) Percentage of hit pairs identified in each family plotted against family size. Color legend indicates the percentage of genes of the respective family involved in hit interactions, either as query or biomarker. j, k) Percentage of hit interactions per gene as a query (j) and as a biomarker (k) is plotted against percentage of hit interactions per family. Color indicates family size.

In summary, this set of proof-of-concept screens – despite their limited scope – demonstrates that *bona fide* paralog dependencies are detectable using pooled LOF screens in cancer cell lines.

### Bioinformatic identification of cancer-relevant candidate paralog dependencies

Due to the limited search space and number of paralog dependencies retrieved from our focused experimental approach, we decided to search for candidate interactions in a systematic manner, leveraging publicly available LOF and expression data from hundreds of cancer cell lines. We hypothesized that candidate genetic interactions between paralogous genes could be detected by looking at the relationship between expression of Paralog A and dependency on Paralog B. In essence, if low expression of Paralog A (“biomarker”) was correlated with sensitivity to loss of Paralog B (“query”), this would identify a potential genetic interaction that could subsequently be evaluated. Depletion scores from complementary datasets^41^ of pooled CRISPR and shRNA screens (AVANA^42^, Sanger^43^ and DRIVE^44^) and corresponding gene and protein expression data^45,46^ for paralog genes were collected. Correlation coefficients and corresponding p-values between expression levels of biomarkers and depletion scores of queries for as many pairs within each paralog family where expression/depletion data were available were then calculated. This approach resulted in a large matrix of correlation tests, containing 14,064 unique genes (biomarkers or queries) and 108,092 unique biomarker-query pairs (Supplementary Table 2), hereafter called PaCT (Paralog Cancer Targets). The obtained correlation coefficients was then filtered via a cutoff of 3*SD (standard deviation) and p-value < 0.05 to generate a hit set (Figure 2a,b; Supplementary Table 2). Simulating the distribution of Spearman correlation coefficients by randomly assigning each query to a gene from different family yields very few interactions at similarly strong correlations, indicating the specificity of PaCT candidate pairs (see Methods; Figure 2c).

In contrast to other recent paralog studies, we did not limit the PaCT search space to paralog pairs by a similarity cutoff or membership in a paralog gene family of a given (small) size^13,19^. An additional advantage of PaCT is the ability to identify candidate interaction partners for genes that have not been targeted themselves as queries, as long as expression data are available for them to act as biomarkers. Overall, of 3,084,147 possible pairs (including self-interactions) from 3,587 paralogue families, PaCT is blind to 2,975,741 pairs (2,795 genes), due to missing depletion and/or expression data for either query or biomarker or both. This also includes pairs where information is available only for a single cell line and, therefore, where no correlation can be calculated. From the remaining 108,406 paralogue pairs, 2,040 unique pairs (1.9%) were identified as significant interactions, and for 106,366 paralogue pairs (98.1%; 14,055 genes), we identified a non-significant correlation in our analysis (Figure 2d).

We then sought to identify possible differences between hit and non-hit paralogue pairs. Insufficient variability in gene/protein expression and/or depletion scores across the cell lines could underlie low correlation coefficients across our dataset. We investigated this for gene expression levels and depletion scores from the AVANA dataset as an example. Indeed, we identified a small, but statistically significant difference in variabilities for both modalities (p-value < 2.2×10^−16^, Kolmogorov-Smirnov test) between hits and non-hits (Figure 2e,f). We further hypothesized that sequence similarity between individual paralog genes could impact the likelihood for an interaction between them. Indeed, based on Ensembl BioMarT DNA sequence similarity, we observed a significant trend that genes involved in significant paralog interactions exhibit higher similarity than those of non-significant pairs (Figure 2g).

Interestingly, significant pairwise candidate interactions are observed between paralogs in families of any size (Figure 2h). Even though many candidate pairs are interactions within 2-member paralog families, we scored significant correlations in larger families of up to 20 members or more. In smaller families, the majority are positive correlations; with increasing family size, the balance shifts towards an even split with negative correlations. On average (AVANA data), we identified 8.4% significant interactions per family when family size is <= 10 that decreased to 1.4% for families containing more than 10 genes (Figure 2i, Supplementary Figure 2 a,b). Finally, we looked at connectivity within paralog families and observed that this varies widely (Figure 2j,k, Supplementary Figure 2c-f). Among the hit pairs, some queries and biomarkers act as hubs, being involved in multiple or all significant interactions, independent of family size. However, other candidates have a more uniform distribution, being identified in only a subset of hit pairs. It remains to be determined what factors underlie these different degrees of connectivity.

As previously described by others^18,47^, some gRNAs in the Sanger and AVANA datasets are promiscuous and match to sites beyond the intended target. For PaCT, we used processed AVANA and Sanger scores ^42,43^ and we confirmed (for the AVANA data as an example) that most genes had zero or one gRNA excluded from the analysis for reasons identified by the investigators of the study (Supplementary Figure 2g; Supplementary Table 2). However, we confirmed previous observations that a sizeable fraction of query genes (25%) had non-uniquely mapping gRNAs assigned to them (Supplementary Figure 2h; Supplementary Table 2), constituting a potential source of false negatives ^18,47^ in our analysis.

Of the 2,472 candidate hit query-biomarker pairs (2,040 of which are non-redundant, involving 2,451 unique genes), 57% displayed a positive correlation between query dependency score and biomarker expression. Most pairs (70%) were found using gene expression data, reflecting the greater robustness of this dataset. We also included genes whose genetic dependency correlates with their own expression, and 20% of our hits are indeed such “self-pairs”. 20 of them have been previously described as CYCLOPS (copy number alterations yielding cancer liabilities owing to partial loss) genes^48,49^ that, when expressed at low levels, are associated with greater sensitivity to further LOF. Similar to previous observations^49^, CYCLOPS genes are overrepresented among the significant hits identified by PaCT (20%), compared to their representation among all potential interactions (13%), mirroring the high frequency of genomic loss in cancer cell lines. In total, our PaCT analysis highlighted 370 unique candidate self-interactions.

The largest proportion of candidate hit query-biomarker pairs (46%) was detected in the AVANA dataset – representing the largest and most comprehensive database in terms of cell lines included -, followed by Sanger (37%) and DRIVE (17%). The covered cell lines and genes overlap to a certain extent, but each dataset contains unique cell lines and genes, in addition to differences in methodologies for generating LOF phenotypes (RNAi in DRIVE vs. CRISPR in AVANA and Sanger)^41^. Thus, it is not surprising that many hits are found uniquely within one dataset or data domain (gene or protein expression; Figure 3a). Nevertheless, 17% (432/2,472) of candidate pairs are recovered more than once, strengthening our confidence in the PaCT approach. The largest overlap was observed between AVANA and Sanger positive correlations pairs using gene expression as a biomarker, confirming those as high-quality candidates. To illustrate the PaCT approach, Figure 3b shows examples of strong negative or positive correlations within small paralog families along the diagonal, i.e. between expression and depletion of the same gene or closely related paralogs, which are listed next to each other.

**Figure 3:**
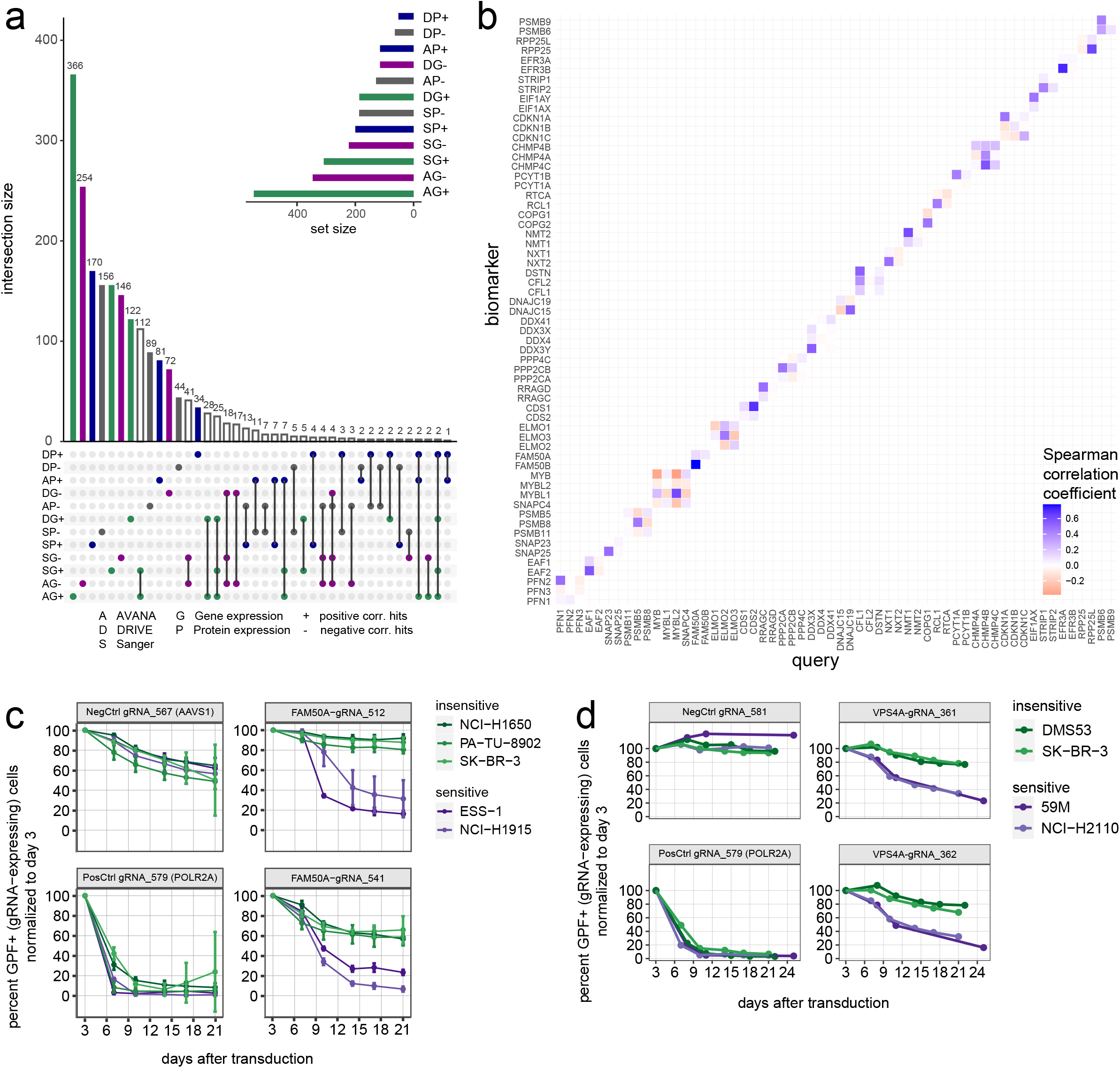
Correlation analysis of public loss-of-function screens yield known and novel candidate paralog genetic interactions. a) Overlap of hit pairs between different input datasets. Y-axis shows the number of overlapping pairs by dataset. Comparisons are indicated by dots and lines on the x-axis, colored by type of expression data (gene, protein) and interaction (pos, neg). Inset shows number of hit pairs by dataset. b) Exemplary pairwise correlation matrix for paralog families of 2-4 members and Spearman correlation > 0.42 for at least one pair in the family including CYCLOPS interactions. c) CRISPR/Cas9 depletion assay in cell lines resistant (green) and sensitive (purple) to loss of *FAM50A*. gRNAs targeting positive control genes (*POLR2A*), negative controls (*AAVS1*) and *FAM50A* are indicated. Cells were lentivirally transduced with the gRNA plasmid containing GFP; GFP percentage in the transduced cell line pool was measured by flow cytometry at the indicated time points and normalized to day 3 post-transduction. d) CRISPR/Cas9 depletion assay in cell lines resistant (green) and sensitive (purple) to loss of VPS4A. gRNAs targeting positive control genes (*POLR2A*), negative controls (*non-targeting*) and *VPS4A* are indicated. Cells were lentivirally transduced with the gRNA plasmid containing GFP; GFP percentage in the transduced cell line pool was measured by flow cytometry at the indicated time points and normalized to day 3 post-transduction.

We complemented the PaCT approach by an additional analysis of the AVANA, DRIVE and Sanger data. First, we classified cell lines as sensitive and resistant to depletion of a given query gene by k-means clustering (k=3, leaving out the intermediate group). Then, we tested whether the expression of a given biomarker gene was significantly different between the sensitive and resistant cluster. The most significant negative correlation hits are almost exclusively self-interactions (Supplementary Figure 3a-c), consistent with the notion of increased sensitivity to loss of highly expressed genes that might act as proliferation drivers. On the other hand, the most significant positive correlation hits are pairs of paralogs (Supplementary Figure 3a-c), supporting the hypothesis of functional redundancy and synthetic lethality between those genes. Overall, the PaCT top hits also emerged as most significant in this analysis.

In order to characterize the PaCT hits, we investigated gene-centric parameters of the candidate pairs (without self-interactions) that have been hypothesized by us and others to affect the likelihood of genetic interaction between paralog genes^13,18^. We observed that some of our candidate interacting paralog pairs (13%) are involved in protein-protein interactions (PPI) with each other, as annotated in BIOGRID^50^ (v4.3.196; Supplementary Figure 3d). We then checked the candidate pairs for homo- and heteromeric interactions^3^, where homomeric means the assembly of a protein with itself whereas heteromeric paralogs assemble with each other. None of the candidate pairs is found on the (short) list of heteromers, and ~3% of queries or biomarkers are annotated as homomers (Supplementary Figure 3d). We also compared our list of candidate pairs to the Critical Paralog Groups (CPGs) defined by Modos et al.^51^, i.e. paralog groups that play important roles in signaling flow and pathway cross-talk. ~2% of PaCT pairs and 4-6% of query or biomarker genes are annotated as members of CPGs (Supplementary Figure 3d). Finally, we investigated whether the candidate interacting pairs share the same common ortholog and whether that ortholog is essential in different model organisms (Supplementary Figure 3e). While these numbers provide a mere estimate, due to the caveats of ambiguous and incomplete ortholog mapping, we expected and observed increasing fractions of common essential orthologs with increasing evolutionary distance – from 0.4% in *M. musculus* to >5.5% in *S. cerevisiae*. Accordingly, the fraction of orthologs that could not be mapped also increased, while the fraction of non-shared orthologs decreased.

Together, these characteristics of shared evolutionary origin and essentiality, or physical interaction, describe some but not all parameters that underlie potential genetic and functional interaction between paralogous genes. Recently, several groups have investigated paralog redundancy and interaction using various computational and experimental methods^13–20^. We compared our PaCT candidates with their sets of potentially interacting paralogs and recovered 12-67% of published pairs in our hit list (Supplementary Figure 3f). Conversely, 15% of PaCT pairs are found in any other dataset. The published sets originate from vastly different search spaces – from a few hundred experimentally tested pairs to computational predictions of the complete interaction matrix of all annotated paralogs. Therefore, the variation in recovery is not surprising and consistent with comparisons between the published datasets^13–20^.

In addition, PaCT also identified several paralog dependencies that have recently been described, including *SMARCA2-SMARCA4* or *SLC25A28-SLC25A37*^19^. We used CRISPR GFP-depletion assays to experimentally validate the genetic dependencies on *FAM50A* in cells where *FAM50B* expression is low, and on *VPS4A* in *VPS4B*-low cell lines (Figure 3c,d and Supplementary Figure 3g,h), two paralog interactions that have recently been functionally characterized^19,52,53^.

While most hits from dual-LOF screens and experimentally validated paralog dependencies rely on the absence of Paralog A to detect dependency on Paralog B (or partial loss in a CYCLOPS interaction), PaCT in principle identifies candidate interactions at any level of expression. To illustrate this, we calculated the fraction of hit pairs with a relevant query depletion (AVANA or Sanger score < −0.5 or DRIVE score < −3) in at least one cell line when the biomarker expression is low, medium or high (Supplementary Figure 3i). Indeed, the theoretical validation rate of candidate interactions is ~60% for all expression bins.

Overall, these findings validate PaCT as a complementary approach to retrieve validated as well as novel candidates for interactions between paralog genes.

### *RPP25-RPP25L* and *DNAJC15-DNAJC19* are novel cancer-relevant paralog interactions

In addition to previously described paralog interactions, we discovered several novel high-confidence candidate dependencies, among them *RPP25-RPP25L*. RPP25 has been described as a component of the RNase P and RNAse MRP ribonuclease complexes that process pre-tRNA and pre-rRNA sequences, respectively^54–57^. Little is known about RPP25L, a role in tRNA or rRNA processing has not been functionally validated. Sensitivity to loss of *RPP25L* was observed to be correlated with low expression of RPP25 (Figure 4a, Supplementary Figure 4a,b). This was then experimentally validated using CRISPR depletion assays. No depletion of *RPP25L*-targeting gRNAs was observed in cell lines that express *RPP25* (Figure 4b,c). Overexpression of either *RPP25* or *RPP25L* in the sensitive U-2OS and KYSE-150 fully rescued sensitivity to *RPP25L* LOF, demonstrating functional redundancy between these two paralogs (Figure 4d, Supplementary Figure 4c). Interestingly, we observed a reduction in levels of endogenous RPP25L upon ectopic overexpression of RPP25 (Supplementary Figure 4d,e) suggesting the existence of feedback mechanisms that regulate the levels of RPP25L in response to changes in the abundance of its paralog protein. In order to elucidate the underlying molecular mechanism of this paralog interaction, further investigation of their role in pre-tRNA and pre-rRNA processing will be required.

**Figure 4:**
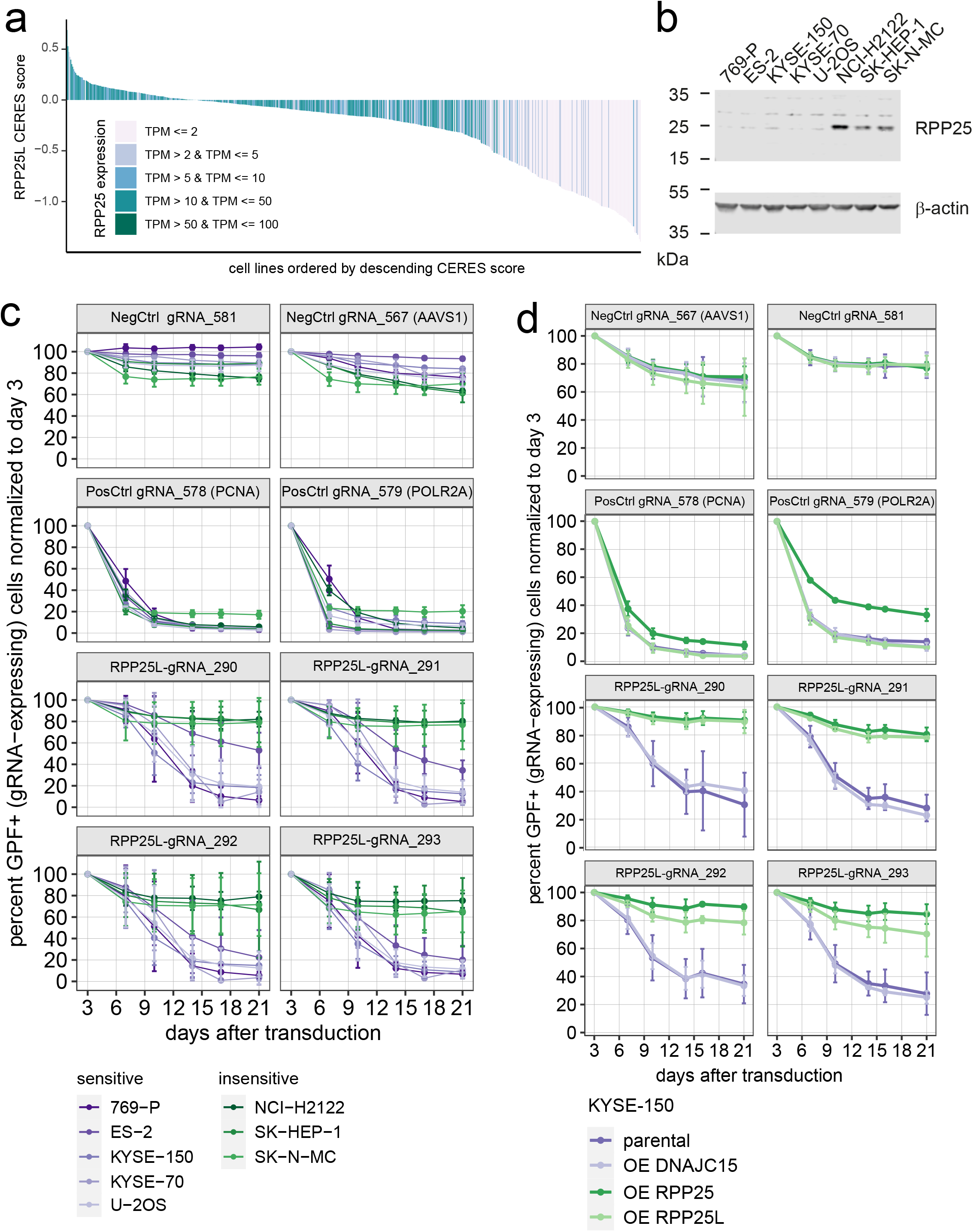
Validation of paralog redundancy between *RPP25* and *RPP25L*. a) AVANA-based depletion scores (CERES) for *RPP25L*, color-coded by *RPP25* expression levels. b) Western blot for *RPP25* in indicated cancer cell lines. β-actin was used as loading control. c) CRISPR/Cas9 depletion assay in cell lines predicted to be sensitive (purple) or resistant (green) to loss of *RPP25L*. gRNAs targeting *RPP25L* (gRNA-290, gRNA-291, gRNA-292, gRNA-293), positive controls (*PCNA, POLR2A*) and negative controls (non-targeting, *AAVS1*) are indicated. Cells were lentivirally transduced with the gRNA plasmid containing GFP; GFP percentage in transduced cell line pool was measured by flow cytometry at the indicated time points and normalized to day 3 post-transduction (n=3 independent replicates of the experiment). d) CRISPR/Cas9 depletion assay as in (c) following ectopic expression of *RPP25* or *RPP25L* in KYSE-150 cells that are sensitive to loss of *RPP25L* (parental). *DNAJC15* expression served as a negative control.

Gene silencing is often accompanied by promoter hypermethylation^58^. We calculated the correlation of methylation levels^46^ of Paralog A with depletion scores of Paralog B and compared the methylation correlation coefficients to the expression correlation coefficients from PaCT. As shown in Figure 5a, (Supplementary igure 5a,b; Supplementary Table 2) for some of the pairs, methylation status of Paralog A could be a useful biomarker for dependency on Paralog B. In particular, we could also detect a negative correlation between methylation and gene expression for multiple CpGs in the promoter regions of *FAM50B* and *DNAJC15* (Figure 5b,c, Supplementary Figure 5c). Although correlation does not necessarily imply causation, it is feasible that methylation could underlie low expression of the biomarker paralog in these cases. DNAJC15-DNAJC19 has not been described as a paralog redundancy before, therefore we set out to validate this interaction experimentally. *DNAJC15* expression levels predict sensitivity of cell lines to loss of its paralog *DNAJC19* according to our PaCT analysis (Figure 5d, Supplementary Figure 5d,e). We confirmed the sensitivity to *DNAJC19* knockout in cell lines that do not express *DNAJC15* (Figure 5e) in CRISPR depletion assays (Figure 5f), including cells with high levels of *DNAJC15* as negative controls. Cell lines that do express *DNAJC15* were predicted to be insensitive to loss of *DNAJC19* and accordingly, *DNAJC19*-targeting gRNAs are not depleted from the pool of cells over time. To conclusively demonstrate functional redundancy between the two paralogs, we overexpressed *DNAJC15i* in the sensitive cell line NCI-H1975 and found that we could thereby rescue the dependency on *DNAJC19* (Figure 5g and h).

**Figure 5:**
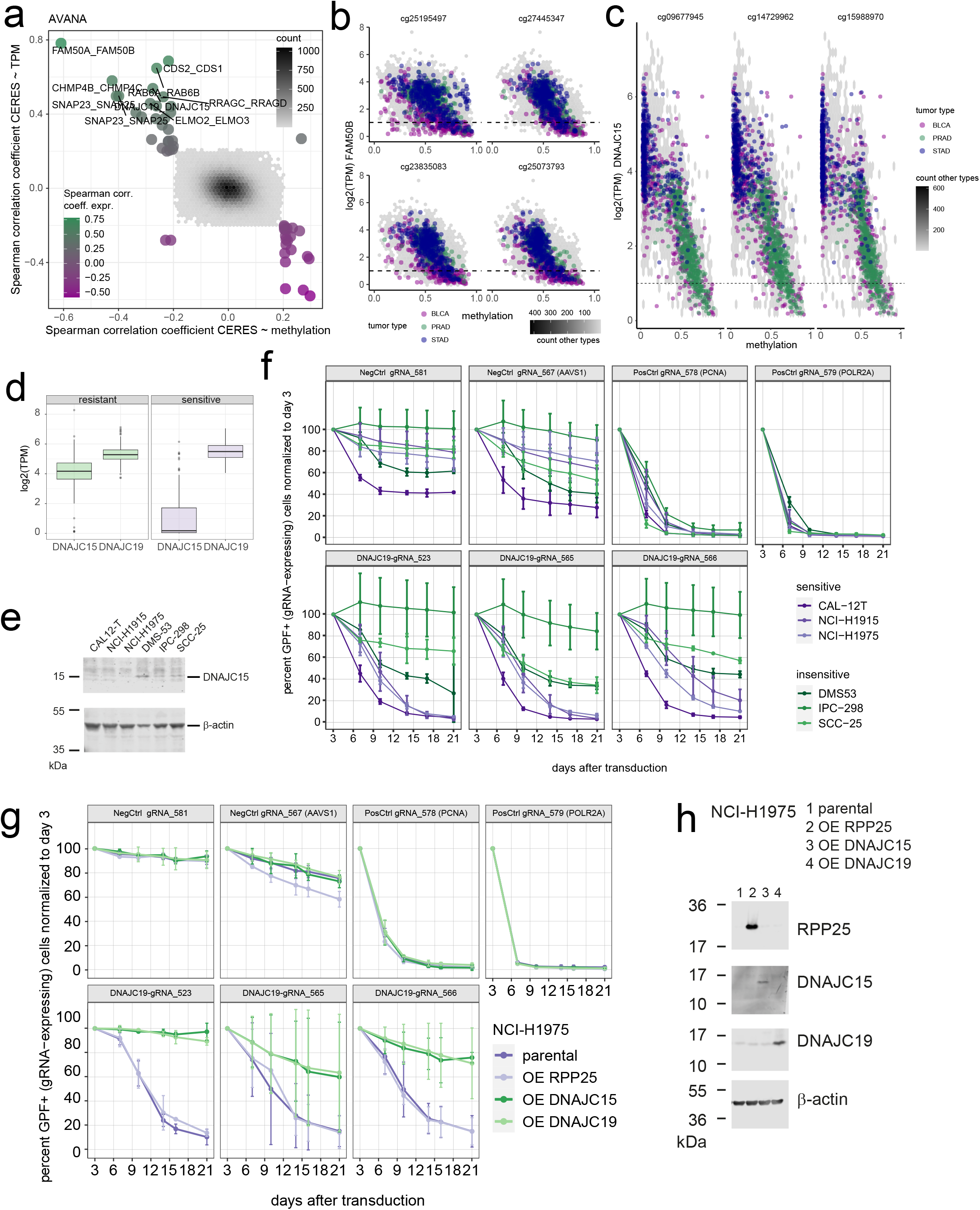
Paralog redundancies for *FAM50A-FAM50B* and *DNAJC15-DNAJC19* can be attributed to expression loss by DNA methylation. a) Scatter plot identifying putative paralog dependencies due to DNA hypermethylation. X-axis: Spearman correlation coefficient between depletion data (CERES score (AVANA data)) and DNA methylation. Y-axis: Spearman correlation coefficient between depletion data (CERES score (AVANA data)) and gene expression (TPM). Pairs with correlation coefficients <|0.2| are displayed as density plots, strongest correlations are labeled. b) Scatter plot of mRNA expression levels (log2(TPM)) of *FAM50B* versus CpG island methylation at indicated loci across tumour types from TCGA. Samples from bladder urothelial carcinoma (BLCA), prostate adenocarcinoma (PRAD) and stomach adenocarcinoma (STAD) studies are highlighted. c) Scatter plot of mRNA expression levels (log2(TPM)) of *DNAJC15* versus CpG island methylation at indicated loci across tumour types from TCGA. Samples from bladder urothelial carcinoma (BLCA), prostate adenocarcinoma (PRAD) and stomach adenocarcinoma (STAD) studies are highlighted. d) Boxplot summarizing expression data (log2(TPM)) for members of the *DNAJC19-DNAJC15* paralog family in cell lines resistant and sensitive to *DNAJC19* loss. e) Western blot of *DNAJC15* levels in selected sensitive (CAL-12T, NCI-H1915, NCI-H1975) and resistant (DMS53, IPC-298, SCC-25) cell lines. β-actin was included as a loading control. f) CRISPR/Cas9 depletion assay in cell lines predicted to be sensitive (purple) or resistant (green) to loss of *DNAJC19*. gRNAs targeting *DNAJC19* (gRNA-318, gRNA-523, gRNA-565, gRNA-566), positive controls (*PCNA, POLR2A*) and negative controls (non-targeting, *AAVS1*) are indicated. Cells were lentivirally transduced with the gRNA plasmids also containing a GFP expression cassette. The percentage of GFP expressing cells in the transduced cell line pool was measured by flow cytometry at the indicated time points and normalized to day 3 post-transduction (n=3 independent replicates of the experiment). g) CRISPR/Cas9 depletion assay in cell lines following ectopic expression of *DNAJC15* in NCI-H1975 cells that are sensitive to loss of *DNAJC19*. Expression was induced by addition of 1 μg/ml doxycycline to the medium at the start of the experiment, which was replenished twice per week. Cells were lentivirally transduced with a gRNA targeting *DNAJC19* (gRNA-318), positive control (*POLR2A*) or negative control (non-targeting). The plasmid also expresses GFP. The percentage of GFP-positive cells in transduced cell line pool was measured by flow cytometry at the indicated time points and normalized to day 3 post-transduction (n=2 independent replicates of the experiment). h) Western blot for RPP25, DNAJC15, and DNAJC19 in NCI-H1975 cells expressing the indicated overexpression constructs upon culture in the presence of doxycycline (1 μg/ml) for 72 hours. β-actin was included as a loading control.

### Paralog buffering between chrX- and chrY-encoded genes

Loss of chromosomes have been reported to be frequently occur during cancer development^21^. Assessing gene expression and copy number data across The Cancer Genome Atlas (TCGA), did not reveal obvious bimodal distributions for any chromosome except chrY, suggesting that that whole chromosome loss is not frequent enough to be detected in this manner across this dataset (Supplementary Figure 6a,b). As described above, LOY has been associated with increasing age and noted in some cancers derived from male patients^21,26–28^. In agreement with this, a bimodal expression distribution for chrY genes within 1.5% of all male TCGA samples, was observed (Figure 6a). Binning samples by tumour purity shows that LOY is more prevalent in samples with higher tumour purity, indicating that LOY could indeed happen more frequently in cancers compared to adjacent normal tissue (Supplementary Figure 6c). Due to the absence of matched non-tumour samples from TCGA, we used data from the Genotype-Tissue Expression (GTEx) project to estimate the frequency of LOY in normal tissues. In corroboration of our hypothesis, at the same 99^th^ percentile cutoff, no LOY was observed across normal samples GTEx (Supplementary Figure 6d).

**Figure 6:**
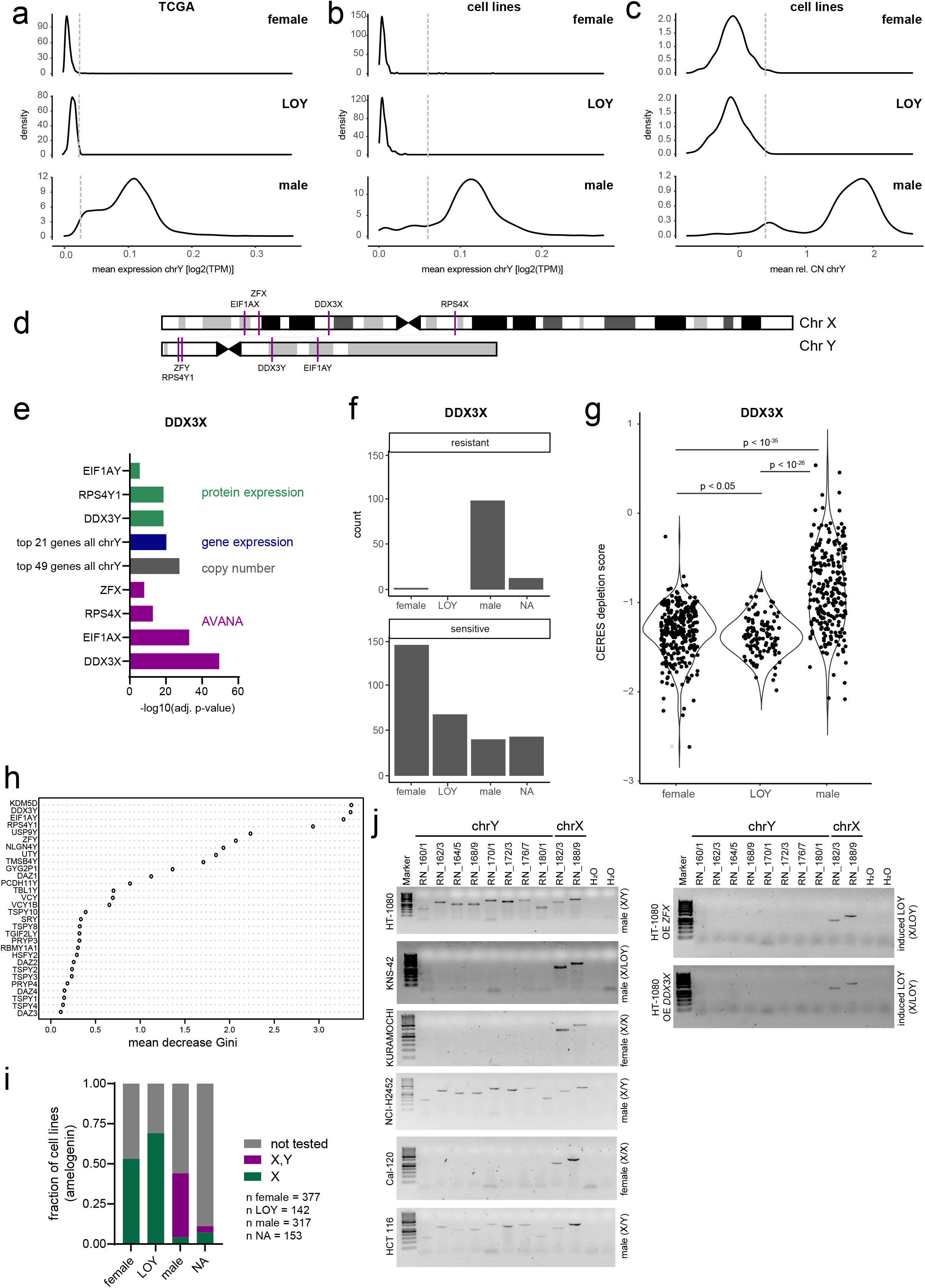
Loss of chrY as potential biomarker for paralog dependencies between sex chromosome genes. a) Distribution of average gene expression (TPM) across genes located on chrY for TCGA samples for which data were available. Sex (male, female) as annotated in TCGA or inferred (LOY) as described in Methods. b) As in (a) for cell lines (CCLE) with available gene expression data. c) As in (b) for average relative copy number (CN). d) Schematic depiction of chrX and chrY with location of interacting paralogs indicated. e) Analysis of factors that are most significantly different between *DDX3X*-loss-sensitive and *DDX3X*-loss-resistant cell lines, as defined using k-means clustering based on AVANA data. For each data domain, the most significant discriminators are displayed. f) Sensitive vs. resistant cell lines (as in (e)) by sex (as in (b and c)). g) *DDX3X* sensitivity (CERES depletion score from AVANA dataset) by sex (as in (b and c)). p-values were calculated using a two-sided Fisher’s exact test for count data with Monte-Carlo-simulated p-value (based on 10000 replicates). h) Variable importance plot for Random Forest model to predict *DDX3X* sensitivity. Gene expression values were used as variables for the Indicated genes on y-axis. i) Fraction of cell lines that harbor chrX and chrY or chrX only, grouped by sex ((as in (b and c)), as assessed by the amelogenin marker in standard STR analysis. j) PCR validation of sex chromosome status in selected cell lines used for further analyses. 8 chrY-specific primer pairs and 2 chrX-specific primer pairs were tested in female patient-derived (KURAMOCHI and Cal-120), male patient-derived chrY retaining (HT-1080 and HCT 116), and male LOY cells (KNS-42).

These studies were further strengthened by analysis of the prevalence of LOY across cancer cell lines used for the AVANA, DRIVE and Sanger screens. LOY was calculated as for the TCGA samples using copy number and expression data and observed in 142 of 459 male cell lines (31% of male, 14% of all cell lines) in our dataset (Figure 6b,c). These studies were supported by analysis of STR profiles for 455 cell lines (46% of cell lines used for PaCT) and analysis of the amelogenin marker for presence or absence of chrY (Figure 6i, Supplementary Table 3)(Figure 6i, Supplementary Table 3). We found that the previous sex assignment was accurate, and LOY status was confirmed for all previously identified cell lines. We further validated the sex chromosome status for a subset of cell lines by a PCR strategy (Figure 6j).

We next investigated whether PaCT retrieved any candidate interactions where the biomarker gene is located on chrY to potentially exploit tumour LOY. Only 24 chrY genes were screened in the AVANA, DRIVE or Sanger datasets, 22 of which are part of our paralog families. Interestingly, in four of these pairs, the query genes are located on chrX: *DDX3X-DDX3Y, RPS4X-RPS4Y1, ZFX-ZFY, EIF1AX-EIF1AY* (Figure 6d). These pairs also rank highly in the predictions by DeKegel et al.^13^. Notably, all four chrX query genes are genes that escape X chromosome inactivation^59,60^, and DDX3X is among a small set of tumour-suppressor genes that escape from X-inactivation (EXITS genes)^61^, where mutations occur more frequently in male cancers and co-occur with LOY.

In order to validate dependency on the chrX paralog when the chrY paralog is not expressed (or chrY is lost), we used CLIFF (*C*ell *L*ine d*IFF*erences)^62^, a web application for the analysis of differences between two sets of cell lines in terms of differential gene or protein expression, DNA copy number, gene signatures, sensitivity to shRNA depletion or CRISPR gene knock-out and other parameters. First, we used k-means clustering to classify cell lines as sensitive and resistant (k=3, leaving out the intermediate group) based on their depletion scores in the AVANA dataset for each of the four chrX paralog hit genes. We then analyzed these groups in CLIFF and looked for the parameters that are most significantly different between the sensitive and resistant cell lines. As a control, we checked that the top gene in the AVANA category is the respective query, i.e. *DDX3X* for the classification run on the *DDX3X* depletion scores (Figure 6e, Supplementary Figure 6e-g). Other AVANA discriminators included some or all of the other chrX hit genes. Conversely, chrY genes, with the respective paralog gene at the top, are the main discriminators based on gene and protein expression, confirming LOY as a potential biomarker that predicts sensitivity to loss of the four selected chrX genes (Figure 6e, Supplementary Figure 6e-g). As expected, LOY cell lines are therefore enriched among the sensitive cell lines for all four chrX genes (Figure 6f, Supplementary Figure 6h-j; p-value sensitive vs. resistant = 10^−4^ for all four genes, Fisher’s exact test). Accordingly, AVANA depletion scores for *DDX3X* (Figure 6g), *EIF1AX, ZFX* and *RPS4X* (Supplementary Figure 6h-m) are generally lower in LOY cell lines than male cell lines. However, some male cell lines are also sensitive to loss of the chrX-encoded paralog, indicating that the genetic buffering by the chrY-encoded gene might be incomplete in some contexts.

Consistent with these analyses, a Random Forest (RF) machine-learning model trained with chrY gene and *DDX3X* paralog family gene expression data on the Sanger depletion dataset predicted sensitive and insensitive cells for the AVANA dataset with an accuracy of 0.82. A variable importance analysis revealed *KDM5D, DDX3Y, EIF1AY* and *RPS4Y1* expression as the top predictors for *DDX3X* sensitivity (Figure 6h). Similar models for *ZFX* and *EIF1AX* were trained on Sanger data, predicted AVANA data with an accuracy of 0.715 and 0.82 respectively (Supplementary Figure 6n,o).

Genetic rescue experiments were performed to validate the putative functional redundancy between chrX/Y-encoded paralogs. *DDX3X* dependency negatively correlates with the expression levels of *DDX3Y* across a panel of >600 cancer cell lines (Figure 7a) i.e. across the AVANA dataset, low expression of *DDX3Y*- but not other family members correlated with sensitivity to *DDX3X* depletion (Supplementary Figure 7a, b). The *DDX3X*-*DDX3Y* functional redundancy was previously suggested in a hamster cell line^63^ and Raji cells^64^ but has not been studied in the context of LOY. In HT-1080 cells, that possess chrY, whereas gRNAs targeting *DDX3X* do not impact the proliferative capacity of these cells, rapid depletion was observed in the context of a gRNA simultaneously targeting *DDX3X* and *DDX3Y* (Figure 7b). Importantly, the effects of the *DDX3X*-*DDX3Y* dual-specific guide could be completely rescued by expression of gRNA-resistant cDNA constructs for *DDX3X* or *DDX3Y*. Similar results were obtained for another male cancer cell line, HCT 116 (Supplementary Figure 7c). KURAMOCHI cells, derived from a female patient, are dependent on *DDX3X* (Supplementary Figure 7d) demonstrating that buffering of the Y-encoded gene is a priori not part of the genetic makeup. Finally, loss of Y-chromosome was confirmed in KNS-42 cells by PCR (Figure 6j). Rapid depletion was observed with gRNAs targeting both *DDX3X* and *DDX3Y* simultaneously as well as gRNA targeting *DDX3X* alone (Figure 7c). Ectopic expression of either *DDX3X* or *DDX3Y* completely reversed the phenotype whereas a functionally unrelated X chromosome located gene X, *ZFX*, did not.

**Figure 7:**
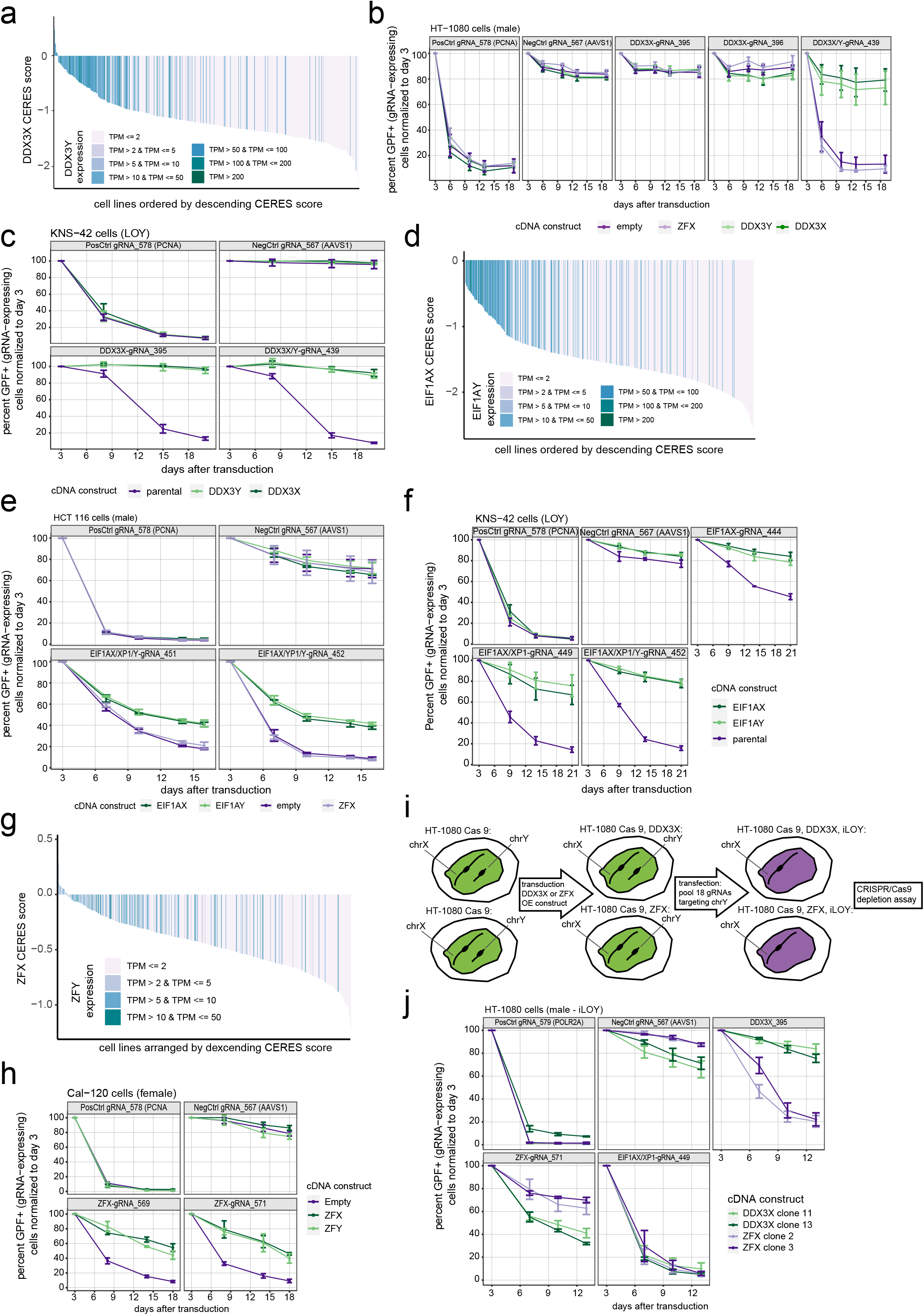
Validation of chrX-chrY paralog dependencies. a) AVANA-based depletion scores (CERES) for *DDX3X*, color-coded by DDX3Y expression levels. b) CRISPR/Cas9 depletion assay in male HT-1080 cells that carry chrY. gRNAs targeting a positive control gene (*PCNA*), negative control locus (*AAVS1*), *DDX3X* (gRNA-395, gRNA-396) and *DDX3X* and *DDX3Y* simultaneously are indicated. Cells were lentivirally transduced with the gRNA plasmid containing GFP; GFP percentage in transduced cell line pool was measured by flow cytometry at the indicated time points and normalized to day 3 post-transduction. Cells were additionally transduced with empty vector (control), unrelated cDNA encoding *ZFX* (control), or rescue constructs with cDNA encoding *DDX3X* or *DDX3Y*. Points in line graph represent mean, and error bars denote the standard deviation (n= 3 independent experiments). c) CRISPR/Cas9 depletion assay in male KNS-42 cells that lost chrY (LOY). Assay, gRNAs and cDNA constructs as in (b). Points in line graph represent mean, and error bars denote the standard deviation (n= 3 independent experiments). d) AVANA-based depletion scores (CERES) for *EIF1AX*, color-coded by EIF1AY expression levels. e) CRISPR/Cas9 depletion assay in male HCT 116 cells that carry chrY. gRNAs targeting positive control (*PCNA*), negative control (*AAVS1*), and *EIF1AX* and *EIF1AY* simultaneously are indicated. Cells were additionally transduced with empty vector (control), unrelated cDNA encoding *ZFX* (control), or rescue constructs with cDNA encoding *EIF1AX* or *EIF1AY*. Assay as in (b), points in line graph represent mean, and error bars denote the standard deviation (n= 3 independent experiments). f) CRISPR/Cas9 depletion assay in male KNS-42 cells that lost chrY (LOY). gRNAs targeting positive control (*PCNA*), negative control (*AAVS1*), *EIF1AX*, *EIF1AX*/*EIF1AXP1*, and *EIF1AX*/*EIF1AXP1* and *EIF1AY* simultaneously are indicated. Cells were additionally transduced with empty vector (control) or rescue constructs with cDNA encoding *EIF1AX* or *EIF1AY*. *EIF1AX*/*XP1* indicates *EIF1AX* and the *EIF1AXP1* pseudogene. Assay as in (b), points in line graph represent mean, and error bars denote the standard deviation (n= 3 independent experiments). g) AVANA-based depletion scores (CERES) for *ZFX*, color-coded by *ZFY* expression levels. h) CRISPR/Cas9 depletion assay in female Cal-120 cells. gRNAs targeting positive control (*PCNA*), negative control (*AAVS1*), and *ZFX* (gRNA-569, gRNA-571) are indicated. Cells were additionally transduced with empty vector (control) or rescue constructs with cDNA encoding *ZFX* or *ZFY*. Assay as in (b), points in line graph represent mean, and error bars denote the standard deviation (n= 3 independent experiments). i) Schematic depiction of workflow for induction of LOY in male HT-1080 cells expressing Cas9 and *DDX3X* or *ZFX*. j) CRISPR/Cas9 depletion assay in male HT-1080 cells where LOY was induced. Two clones each expressing cDNA constructs encoding *DDX3X* or *ZFX* were transduced with gRNAs targeting positive control (*POLR2A*), negative control (*AAVS1*), *DDX3X*, *ZFX* or *EIF1AX*/*EIF1AXP1*. *EIF1AX/XP1* indicates *EIF1AX* and the *EIF1AXP1* pseudogene. Assay as in (b), points in line graph represent mean, and error bars denote the standard deviation (n= 3 independent experiments).

These findings were then extended to additional PaCT genes with a putative chrX/Y-encoded redundancy. Sensitivity to *EIF1AX* correlates with the expression of *EIF1AY*, similar to *DDX3X-DDX3Y*, (Figure 7d, Supplementary Figure 7e, f). LOY resulted in a strong dependency on *EIF1AX* (Figure 7f) whereas cells retaining chrY were only sensitive to gRNAs simultaneously targeting *EIF1AX* and *EIF1AY* (Figure 7e). Depletion could be reversed upon expression of gRNA-resistant cDNA constructs encoding for *EIF1AX* or *EIF1AY* (Figure 7e, f). Similar results were obtained with gRNAs targeting *RPS4X* or *RPS4Y*, (Supplementary Figure 7g). In addition to *DDX3X-DDX3Y* and *EIF1AX-EIF1AY, ZFX-ZFY* emerged as an additional functionally redundant paralog pair from our PaCT analysis. Sensitivity of cancer cell lines to the loss of *ZFX* correlates with the expression of *ZFY* and, less strongly, with *ZNF711* (Figure 7g, Supplementary Figure 7h,j). As the sensitivity to *ZFX* loss-of-function is less pronounced in KNS-42 cells in the AVANA dataset^42^, we turned to female Cal-120 cells for depletion and rescue experiments. CRISPR/Cas9-mediated loss of ZFX resulted in depletion of GFP- and gRNA-expressing cells. This phenotype could be rescued with gRNA-resistant cDNA constructs encoding for *ZFX* or *ZFY*, validating the functional redundancy between the two proteins (Figure 7h).

In order to confirm that loss of chrY is the causative event in dependency on the paralogs encoded on chrX we designed an approach to engineer removal of chrY (induced LOY, iLOY). Similar to published approaches that have demonstrated loss of the targeted chromosome ^65,66^, a pool of 18 gRNAs targeting chrY genes was introduced in HT-1080 cells ectopically expressing either *DDX3X* or *ZFX* ^62,63^ (Figure 7i). LOY was validated by PCR (Figure 6j). Two independent clones were derived and subsequently treated with gRNAs targeting *DDX3X*, *ZFX* and *EIF1AX*. Whereas no phenotype was observed in parental HT-1080 cells (Figure 7b,f), LOY clones were sensitive to gRNAs for the chrX-encoded paralog genes. This sensitivity was lost or reduced upon ectopic expression of corresponding gRNA resistant constructs, e.g HT-1080 iLOY *ZFX* are sensitive to a gRNA targeting *DDX3X* whereas HT-1080 iLOY *DDX3X* are not (Figure 7j and Supplementary Figure 7j).

Altogether, these data suggest that selective targeting of paralogs encoded on the X-chromosome, for which genetic buffering with a chrY-encoded gene exists, might be a generalizable strategy to target LOY tumours. The iLOY experiments validate the loss of chrY as the root cause for this dependency. To the best of our knowledge, these are the first examples for synthetic lethal interactions between paralogs located on the X and Y chromosomes.

## Discussion

Exploiting distorted genetic buffering in human malignancies represents a promising therapeutic concept. The clinical activity of poly ADP ribose polymerase (PARP) inhibitors in cancers with defects in the homologous recombination-based DNA damage repair pathway^67–69^ underlines this point. Paralog genes, originating from gene duplication events, represent an additional subset of these general synthetic lethal genetic interactions where tumour-specific loss of a paralog gene creates a therapeutically exploitable dependency on the remaining paralog gene. In this study, we identified >2000 candidate paralog dependencies relevant to human cancer. We have experimentally validated a subset of these paralog pairs and provide evidence that genetic buffering between the sex chromosomes could provide an attractive therapeutic strategy for human cancers of individuals that have lost the Y chromosome in malignant cells.

Our analysis was confined to cancer-relevant interactions that can be identified in the respective cell lines used and genes targeted in publicly available CRISPR/RNAi LOF screens. Due to lack of equal representation of different cancers within the datasets this could lead to a bias for certain tumour types. As described, our discovery pipeline is also “blind” to certain other cases, including uniform expression or depletion of a paralog across all screened cell lines. This is because expression-dependency calculations rely on varying gene expression and depletion scores of one paralog gene across these cell lines. Therefore, approaches like PaCT together with combinatorial genetic screens will further advance our understanding of genetic redundancies. It will be interesting to determine if paralog interactions can be tissue specific and if, within larger families, subsets of genes can have a greater or lesser functional redundancy – a result suggested by our study. If true, this could hint towards the resistance of sub-families and help to functionally annotate understudied paralog genes.

As the PaCT approach relies on publicly available screening data, the caveats of the original experiments, such as suboptimal gRNA design in some instances, are carried over into our dataset. The *DNAJC15-DNAJC19* example illustrates such a case, where all gRNAs in the public dataset also target a pseudogene sequence. While our experimental validation uses independently designed gRNAs, a potential partial function of the presumed pseudogene will have to be determined. Furthermore, additional investigations will show whether *DNAJC15* and *DNAJC19* indeed both play a role in mitochondrial morphogenesis, and whether *RPP25L* is a bona fide subunit of the RNase P/MRP complexes.

A number of mechanisms can underlie the paralog loss. In addition to mutation and deletion we provide evidence that epigenetic mechanisms can also play a role. Validated paralog pairs *DNAJC15-DNAJC19* and *FAM50A-FAM50B*, provide examples where high promoter methylation could, in part, account for decreased expression of one paralog gene. This suggests that DNA hypermethylation in tumours could expose novel vulnerabilities that could be exploited therapeutically. Future research will have to clarify if vulnerabilities originating from DNA hypermethylation are stable enough to permit long-term treatment.

Our study revealed extensive genetic redundancy between the sex chromosomes. We identified four candidate paralog dependencies (*EIF1AX-EIF1AY*, *DDX3X-DDX3Y*, *RPS4AX-RPS4Y1* and *ZFX-ZFY*) of which we validated three experimentally. Our data suggest that cell lines originating from individuals with chrX and chrY become sensitive to the loss of the chrX-encoded gene upon loss of chrY. While this concept could in principle be exploited therapeutically to treat LOY tumours, premalignant states of mosaic LOY in hematopoiesis or ageing-associated LOY, several hurdles would have to be overcome. It would be important to ensure selectivity of the targeting therapeutic between highly similar paralogs. Although we have not observed LOY across the GTEx dataset, it is possible that alternative mechanisms may also lead to down-regulation of the chrY expressed paralog in normal tissues. While not explicitly addressed, recent studies imply incomplete redundancy for *EIF1AX-EIF1AY* and *DDX3X-DDX3Y* in different contexts in absence of LOY^70–72^.

Overall, our study identifies cancer-relevant paralog dependencies and provides a framework for validation and future discovery as further panels of functionally validated cancer cell lines become available. While our PaCT approach currently addresses gene expression, deletion and methylation in the paralog genetic space, the approach is generalizable and could be performed analogously for non-paralog genes as queries, and mutations, passenger deletions or other tractable aberrations as biomarkers. We envisage that this will identify additional testable hypotheses for targeted cancer treatment.

## Materials and Methods

### Cell culture

All cell lines and the respective media are listed in Supplementary Table 4. Cell lines were regularly checked for mycoplasma, authenticated by STR profiling (Eurofins Genomics) and kept at low passage numbers in humidified incubators at 37°C and 5% CO_2_.

### Generation of Cas9- and paralog-expressing cell lines

cDNA sequences for Cas9 and paralog genes were human codon-optimized, synthesized and cloned into their respective vector backbone (Supplementary Table 4) at Genscript Biotech Corporation. Cells were lentivirally transduced. Viral particles were generated using the Lenti-X Single Shot System (Clontech). 72 hours later, stable transgenic cell pools were selected using puromycin or blasticidin (see Supplementary Table 4 for details).

### CRISPR/Cas9 library design, cloning and virus production

The majority of genes in the gRNA library were manually selected from (i) paralog families of 2-5 members, (ii) genes frequently deleted in TCGA samples with a focus on deep deletions in lung adenocarcinoma, lung squamous cell carcinoma, colon adenocarcinoma, liver hepatocellular carcinoma, pancreatic adenocarcinoma, ovarian serous cystadenocarcinoma and prostate adenocarcinoma. gRNA sequences were selected to target protein domains (annotated using PFAM domain identifiers) as described^34^, as well as control sequences for a total of 9574 gRNAs (Supplementary Table 1).

Pooled gRNA oligonucelotides (20-mer target sequences plus cloning adapters; TGCTGTTGACAGTGAGCGCGTCTCTCACCG[20×N]GTTTGGAGACGCCTAGGATCGACGCGGACAACA; Twist Bioscience) were PCR-amplified (0.1 ng DNA input, 24 parallel reactions, 15 cycles). Pooled reactions were purified using the QIAquick PCR purification kit (Qiagen) and digested with BsmBI. The vector backbone (lentiviral vector coexpressing sgRNA, GFP and NeoR, similar to sgETN^73^) was prepared by BsmBI digestion, dephosphorylation and purification as above. Ligation was performed in 14 parallel reactions using T7 ligase and remaining uncut backbone was removed by BsmBI digestion. Ligation products were purified by phenol extraction, transformed into MegaX DH10B T1 electrocompetent bacterial cells (Invitrogen) following manufacturer’s protocol and plated on LB/Ampicillin plates. Colonies were combined and maxi-preps were performed at ~7000× colonies per sgRNA.

Lentivirus was produced in 293T-Lenti-X cells (Clontech) using 10 μg of library DNA and Ready-to-use Lentiviral Packaging Plasmid Mix (Cellecta, 0.5 μg/μL) per 10 cm dish (20 dishes in total). 293T-Lenti-X were plated without antibiotics and transfected the next day using Lipofectamine LTX & Plus (Thermo Fisher). Medium was changed after 7 h of incubation and viral supernatant was harvested after 48 h. Virus titration was carried out individually for each cell line using three different amounts of viral supernatant in the presence of 8 μg/mL polybrene. Transduction efficacy was evaluated 72 h after infection by measuring GFP expression by flow cytometry.

Primer sequences are listed in Supplementary Table 4.

### CRISPR/Cas9 screens

Cas9-expressing cell lines were transduced with the sgRNA library at a multiplicity of infection of ~0.3 in the presence of 8 μg/mL polybrene. To this end, 44 × 10^6^ cells were cultured in four or more T175 flasks for 12/18 population doublings, representing 1000-fold library coverage. Cell numbers were adapted according to measured GFP percentage after initial infection. From a pellet of the respective cell number at the end point, genomic DNA was isolated using the QIAamp DNA Mini Kit (Qiagen). Amplicons around the sgRNA sequences were PCR amplified (1 μg input per PCR reaction, 29 cycles) with barcoded primers. The total amount of genomic DNA input was calculated by dividing the used total cell number by the assumed value of 6 pg genomic DNA per cell. PCR products were purified using the QIAquick PCR purification kit (Qiagen) and a 2% agarose gel using the QIAquick gel extraction kit (Qiagen). In a second PCR, 10 ng of the purified product per reaction were amplified (5 cycles). The pooled PCR products were purified using the QIAquick PCR purification kit. 50 ng of amplicons were used for the library generation with the TruSeq Nano DNA Library Prep kit for NeoPrep (Illumina). The sequencing was conducted on a HiSeq1500 (Illumina) in rapid mode with the paired end protocol for 50 cycles. For the 7 cell lines (MIA PaCa-2, Hep 3B2.1-7, NCI-H1373, NCI-H1993, NCI-H2009, PC-9, HuP-T4) total read counts ranging from 3.1M to 41.6M were generated. Primer sequences are listed in Supplementary Table 4.

### CRISPR/Cas9 library quality control and screen analysis

For the plasmid library, 20 million reads were generated and the gRNA representation was tested for uniformity. gRNA counts ranged from 50 to 8708 reads (25^th^ percentile: 983; median: 1682; 75^th^ percentile 2560 reads). For screen analysis, we used the ‘mageck test’ function of the MAGECK tool (version 0.5.6)^74^ to determine the log2-fold-changes and significance estimates (p-values, FDR) for gRNA representation differences between any of the 7 cell lines and those observed in the plasmid library using the following parameters: “mageck test --norm-method control --gene-lfc-method median”.

To further assess the technical quality of the screens, we overlapped the library with known core-essential (n=625) and never-essential (n=1344) genes constructed from genome scale screens. We found that 307 and 596 gRNAs targeted a subset of the core- and never-essential genes, respectively. We observed a good separation of both guide sets (strictly standardized mean difference < −0.9) and a strong enrichment of core-essential genes in the top depleted genes (AUC > 0.9). Both quality metrics were calculated based on log2-fold-changes from the comparison to the gRNA representation in the plasmid library.

To compensate for the variable effect sizes from the different cell lines, we scaled all gene-level log2-fold-changes such that the median log2-fold-change of all never-essential and core-essential genes where set to 0 and −1, respectively. We call this scaled log2-fold-change escore (essentiality score).

For hit calling, we selected genes that were specifically depleted (cutoffs for escore < −0.4 and FDR < 0.1) in cell lines that harbor a deletion of a member of the same paralog family (absolute copy number = 0 and log2 relative copy number < −1).

### TCGA data

For gene expression data, the GDC Data Portal’s interface (https://portal.gdc.cancer.gov/) was used to compile all data files that mapped the fields “Program” = “TCGA”, “Data Type” = “Aligned Reads”, “Experimental Strategy” = “RNA-Seq”, and “Workflow Type” = “STAR 2-Pass”. Using the GDC Data Transfer Tool, the data was transferred and pre-processed using samtools^75^ collate and fastq to generate FASTQ files, containing the unmapped reads. All samples were subsequently processed with a harmonized RNA-seq pipeline^76^.

TCGA SNP6 copy number segmentation data was downloaded from NIH GDC (https://portal.gdc.cancer.gov/) on December 3 2018. The segmentation information was obtained from the files *nocnv\_grch38.seg.v2.txt. Gene-wise copy numbers were determined by overlapping the segmentation information with Ensembl v86 gene annotation. If a gene was covered by a single segment, the copy number of the segment was assigned to the gene. If a gene was covered by multiple segments, a weighted average copy number was computed based on the size of the overlap between the gene and each segment. Relative copy numbers <= 1.0 were considered as “deep deletion”. The R package TCGAbiolinks (v2.5.9)^77^ was used to extract sample and patient information for TCGA samples by using a custom-made R script.

The sample cohorts COADREAD, FPPP, GBMLGG, KIPAN, and STES were excluded. Data for TCGA methylation loci plots were downloaded from http://www.bioinfo-zs.com/smartapp/^78^. Gene expression levels (log2(TPM)) were plotted against methylation levels of CpGs belonging to islands located in promoter regions of genes of interest.

### Cancer Cell Line Encyclopedia (CCLE) data

Cell line names and descriptions (including sex) were taken from the provider’s cell-line data sheet. If a cell line was available from various vendors, the cell-line name was taken from the top rank in a hierarchy of vendors in the following order: ATCC, DSMZ, ECACC, JCRB, ICLC, RIKEN, KCLB. For gene expression, raw FASTQ data for all CCLE cell lines^46^ were downloaded via the European Nucleotide Archive (accession number PRJNA523380). All data were processed identically to TCGA data as described above.

For copy number determination, SNP6 CEL files were downloaded from https://cghub.ucsc.edu/ in October 2012. Relative copy number segments were computed using the R packages aroma.affymetrix (v3.1.0)^79^ and Rawcopy (v1.1)^80^: SNP6 data were processed with the AROMA method CRMA v2, where the 50 samples with the least amount of copy number alterations based on Rawcopy were used to calculate the reference intensities. This was followed by CBS segmentation. Afterwards, the copy number segments were overlapped with Ensembl v86 gene annotation as described for the TCGA data in order to obtain gene-wise relative copy number values. “Deep deletion” status was assigned as for TCGA data. Absolute copy number segments were computed using PICNIC version c_release 2010-10-29^81^ with reference files adapted for reference genome hg38 and default parameters. The resulting segments were overlapped with Ensembl v86 gene annotation as for TCGA data in order to obtain gene-wise absolute copy number values.

Methylation^46^ data are ‘CCLE_RRBS_TSS1kb_20181022.txt.gz’, downloaded from https://portals.broadinstitute.org/ccle/data. Protein expression^45^ data were directly exported from the indicated reference.

### GTEx data

GTEx v8 gene expression data (phs000424.v8) where processed as described above (RNA-seq pipeline v2.0 (C-GET)^76^). For 4 samples processing failed, and 582 samples failed QC based on sequence length, GC content, assigned reads, intronic bases, 3’/5’ biases, uniquely mapped reads or *GAPDH* detection, and were not included into the final object. Samples from the “Cells – Transformed fibroblast”, “Cells – EBV-transformed lymphocytes” and “Cells – Leukemia cell line (CML)” classes are omitted from the data set.

### CRISPR/Cas9 depletion assays

All CRISPR/Cas9 depletion assays were conducted as previously described^82^. In brief, gRNA sequences were cloned into their respective vector backbone, typically containing GFP (Supplementary Table 4), at Genscript Biotech Corporation. Lentiviral particles were produced in 293T-Lenti-X (Clontech) cells cultured in DMEM, 10% Tet-system approved FCS, 1X Glutamax, 1X NaPyr. 4 × 10^6^ cells were plated in 8 ml medium in 10 cm dishes and transiently transfected with 7 μg of plasmid DNA mixed with Lenti-X Packaging Single Shots (VSV-G) (TakaraBio) according to the manufacturer’s instructions on the following day. 4 hours after transfection, 6 ml fresh medium was added to the plates. Supernatant was harvested 48 hours after transfection, filtered through a 0.45 μm PVDF filter (Millipore) and stored at −80°C in unconcentrated aliquots until further use. Relevant cell lines stably expressing Cas9 (see Supplementary Table 4) were plated at approximately 50 –60 % confluence in 12 or 24 well plates and transduced with 250-500 μl of gRNA virus to achieve 10%-95% transduction efficiency. After transduction, the fraction of GFP positive cells was determined at indicated timepoints using flow cytometry.

Where cell lines expressing doxycycline-inducible cDNA constructs were included in depletion assays, expression was induced at the start of the experiment by addition of 0.5-1 μg/ml doxycycline to the medium, which was thereafter replenished twice per week.

### siRNA assay

Cells were seeded at a density of 4 × 10^5^ in 6-well plates in standard culture media. 24 hours after seeding, cells were transfected with OTP Smartpool reagents (Horizon Discovery) targeting *CSTF2* individually or in an equimolar mixture, *CSTF2T* or negative control at a final concentration of 20 nM using RNAiMAX (Invitrogen) as specified by the manufacturer. 24 hours post transfection media was exchanged and cells further incubated for 48 hours. siRNA details are listed in Supplementary Table 4.

### cDNA overexpression

Constructs based on the pMSCV-Linker-PGK-Blasti backbone (see Supplementary Table 4) were packaged into viral particles using the Platinum-GP Retrovial Packaging Cell line (). Briefly, 5 × 10^6^ cells were plated in 10 cm dishes and co-transfected with 3 μg VSV-G plasmid and 9 μg of the respective construct Lipofectamine LTX (Thermo Fisher) the following day. Medium was exchanged after 16 h and harvested 48 h later for filtration using 0.45 μm PVDF filter (Millipore) and subsequent storage at −80 °C before transduction of target cells and subsequent selection of successfully transduced cells through addition of Blasticidin to the medium.

Constructs based on the RT3REN backbone (see Supplementary Table 4) were packaged into lentiviral particles using the Platinum-E packaging cell line (Cell Biolabs). In brief, 600,000 cells were plated in 6 well plates and transfected with 2 μg plasmid DNA using 6 μl Lipofectamine LTX reagent (Thermo Fisher). Medium was exchanged after 16 h and harvested 24 h later, filtered through a 0.45 μm PVDF filter (Millipore) and added directly to recipient cells stably expressing an ecotropic receptor (pRRL-RIEH), followed by selection with Geneticin. Lentivirus for pRRL-RIEH was produced in lenti X 293T-Lenti-X (Clontech). 4 × 10^6^ cells were plated in 8 ml medium in 10 cm dishes and transiently transfected with 7 μg of plasmid DNA mixed with Lenti-X Packaging Single Shots (VSV-G) (TakaraBio) according to the manufacturer’s instructions on the following day. 4 hours after transfection, 6 ml fresh medium was added to the plates. Supernatant was harvested 48 hours after transfection, filtered through a 0.45 μm PVDF filter (Millipore) before addition to cells and subsequent selection with Hygromycin.

### Western blot

Cells were lysed using RIPA buffer (Sigma) supplemented with HALT protease and phosphatase inhibitor cocktail (Thermo Fisher). Lysates were incubated on ice for 30 min, centrifuged at 14,000 rcf for 20 min at 4°C and protein amounts in the supernatant determined using the Bradford assay (BioRad) according to the manufacturer’s instructions. Laemmli buffer was added to samples followed by boiling at 95 °C for 5 min. Samples were loaded on a pre-cast gel (Criterion XT Precast 4-12 % Bis-Tris Gel, BioRad), run at 150 V for 1.5 hours in XT MOPS running buffer (BioRad) before transfer onto a nitrocellulose membrane (Transblot Turbo Transfer Pack Midi 0.2 μm) for 7 min using the Transblot Turbo Transfer System (BioRad, program: Quickblot Mixed MW, Midi Gel). Membranes were incubated for 1 hour in blocking buffer (10% BSA, 10% PBS-T in water) followed by overnight incubation at 4 °C with primary antibody in BSA antibody buffer (5 % BSA in PBS-T). The next day, membranes were washed three times with PBS-T (10 min per wash) and incubated with secondary antibody in Casein antibody buffer (0,1% Casein in PBS-T) for 1 hour in the dark at room temperature. Membranes were washed three times in PBS-T (10 min per wash) and visualized on an Odyssey CLx imaging system (LI-COR Biosciences).

All antibody details can be found in Supplementary Table 4.

### Correlation analysis (PaCT)

Depletion data for individual genes were obtained from three studies: DRIVE^44^ (2017-10-01), AVANA^42^ (21Q1) and Sanger^43^ (Release 1). Subsequently, depletion values for every screened gene with unique gene symbols were correlated to expression values (TPM, see above), methylation^46^ or protein expression^45^ data across the screened cell lines. Methylation data were summarized for genomic regions mapping to a gene. Pearson, Spearman and Kendall correlation coefficients and corresponding p-values were collected. The gene with depletion data is referred to as *query (q)* gene and the gene with expression/methylation data is referred to as *biomarker (b)* for pairwise correlations. Subsequently, data were filtered for genes which are part of a paralog family, such that every pairwise correlation between *q* and *b* is considered if *q* and *b* are part of the same paralog family:

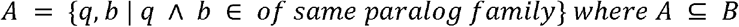

where *B* denotes all correlations between *query* and *biomarker* pairs (screened genes (*q*) and genes with protein and/or mRNA expression values(*b*)) and *A* denotes all correlations for a given paralog family.

We used Spearman coefficients and p-values for all subsequent analyses to account for possible non-normal distributions in the data and minimize the impact of outlier values. Due to differences in query and cell line libraries used, and different scoring systems, each sub-dataset that was processed separately (AVANA, Sanger and DRIVE scores for gene and protein expression). For each sub-dataset, we calculated its own cutoff at 3*SD (standard deviation) and additionally filtered for p-value < 0.05. For gene expression data, all p-values at the 3*SD cutoff were highly significant, likely due to the more complete source data for this domain.

The complete PaCT results can be found in Supplementary Table 2.

### PaCT exploratory space

For all 3,587 paralog families with at least two members, we computed all possible pairwise interactions across members of the same paralog family, including self-interactions. This approach resulted in a total of 3,0841,147 potential pairs. We then assessed the potential of our approach to detect and quantify pairwise dependencies by depletion-expression correlation. Pairs where the query gene was not targeted in any depletion dataset and/or targeted in zero cell lines, and without gene expression data for the biomarker gene expression were labeled as “no info both”. Pairs for which information was missing for either depletion or gene expression were labeled as “no info query” and “no info biomarker”, respectively. Pairs for which information was available in only a single cell line do not allow calculation of a correlation and were labeled “1 cell line”. Pairs for which information was available both for the query and biomarker in at least two cell lines was labeled as “info query & biomarker”. Protein expression data were not included in this analysis.

### PaCT simulation analysis

To identify the difference between PaCT hit correlations and random correlations, we performed 1000 simulations for each gene from each family with significant interactions. For each query gene *q* from each family *f*, we generated a vector of genes *v* with same size as *f*. The new set of genes in *v* contains only the query gene from *f* but the remaining genes in *v* are sampled without replacement from the remaining paralogue families. Then pairwise correlations were computed as above.

### Wilcoxon-test analysis

Cell lines were split into sensitive, resistant and intermediate groups using a k-means clustering algorithm with k=3 for the depletion scores of every gene in the DRIVE^44^, AVANA^42^ and Sanger^43^ datasets. For cell lines in the sensitive and resistant bins, gene and protein expression data were collected. Subsequently, a non-parametric test (Wilcoxon test) was conducted for all query-biomarker pairs. p-values were collected and corrected for multiple testing (Benjamini-Hochberg). The query-biomarker pairs were then filtered as described above.

### Random Forest model

CCLE gene expression data of chrY-encoded and *DDX3X, EIF1AX, ZFX* and *RPS4X* paralog family genes were used to train a Random Forest (RF) model on the Sanger^43^ depletion data. k-means clustering (as described above) was used to separate sensitive and insensitive cell lines. Subsequently, AVANA^42^ sensitivity data were predicted using the RF model.

### Ortholog analysis

We first converted our list of human paralogs to the best match ortholog in *M. musculus, D. melanogaster, D. rerio, C. elegans* or *S. cerevisiae* using DIOPT^83^ (v8). Then we added information on each gene’s essentiality from OGEE^84^ (v2), with a majority vote decision on calling a gene essential or non-essential in cases where more than one dataset with ambiguous calls existed. For all paralog interaction hit pairs, we then checked whether query and biomarker share the same ortholog gene and if so, whether the ortholog is essential.

### Multi-mapping gRNA analysis

We downloaded information on dropped gRNAs and gRNA mapping for the AVANA library from https://depmap.org/portal/download/. Based on this information, we extracted the number of dropped, uniquely mapped or multi-mapping gRNAs for each query gene in the list of PaCT pairs.

### LOY inference

In addition to gene expression and copy number (CN) data, TCGA, GTEx and CCLE provide annotation of the sex of the patients where a sample/cell line originated from. We calculated (i) the average TPM, (ii) the maximum TPM, (iii) the average raw count, (iv) the average relative CN, and (v) the average absolute CN for all genes located on chrY for all samples. TCGA and GTEx do not provide CN data for chrY. For samples originally annotated as male, we checked whether *all* of their values (i)-(v) for cell lines and (i)-(iii) for tissue samples were below the respective 99^th^ percentile of female samples. If this was the case, we re-annotated the sample as LOY.

### PCR validation of LOY

Genotyping primer pairs for different genes on chrX and chrY were designed and tested for specificity. Genomic DNA was extracted from female, male and LOY cells using the QIAamp DNA Mini Kit (Qiagen) following the manufacturer’s protocol. PCR was run using AmpliTaqGold DNA polymerase (Thermo Fisher Scientific) with 100 ng genomic DNA as input. 55°C annealing temperature was used for all primer pairs. Resulting amplicons analyzed on a 2% agarose gel. All primer sequences are listed in Supplementary Table 4.

### Induction of LOY

One million HT-1080 cells expressing Cas9 (puromycin) and *DDX3X* (blasticidin) or *ZFX* (blasticidin) constructs were transiently transfected with a pool of 18 GFP-containing plasmids encoding for gRNAs targeting different chrY genes (RN-gRNA_429-434, RN-gRNA_441-443, RN-gRNA_458-466 using Lipofectamine 3000 (Thermo Fisher Scientific) according to the manufacturer’s instructions. 48 hours after transfection, GFP-positive cells were isolated by FACS and diluted to obtain single cell clones. Clones were screened for LOY by PCR from genomic DNA (as described above) using standard laboratory techniques. Clones with PCR products for chrX but without PCR products for chrY were selected. gRNA sequences are listed in Supplementary Table 4.

### Software and data availability

All calculations were performed in R. Data were visualized using R or GraphPad Prism. All data are publicly available through the indicated references and provided as Supplementary Material, including an R Markdown script containing all code and versioning information to reproduce analyses and figures.

## Acknowledgements

We wish to thank Norbert Schweifer, Tamara Tröls, Harald Studensky and Silvia Blaha-Ostermann for technical assistance, Johannes Zuber for help with cloning the paralog gRNA library, and all colleagues at Boehringer Ingelheim RCV Cancer Research Target Discovery for discussions and critical input to the manuscript.

## Author Contributions

A.K., A.H., F.S., T.P., S.O., C.W., M.C., and C.R. conducted wet lab experiments. A.K., A.S., A.P., V.T., F.S., B.M. and R.A.N. conducted bioinformatic analyses. R.A.N. and B.M. conceived study. B.M., R.A.N. and M.P. oversaw study. J.P., S.W., A.S., led paralog library screens. A.S. helped conceptualize study. A.K., B.M. and R.A.N. wrote manuscript with input from all other

## References

1. Ihmels, J., Collins, S. R., Schuldiner, M., Krogan, N. J. & Weissman, J. S. Backup without redundancy: genetic interactions reveal the cost of duplicate gene loss. Mol Syst Biol 3, 86 (2007).

2. Vavouri, T., Semple, J. I. & Lehner, B. Widespread conservation of genetic redundancy during a billion years of eukaryotic evolution. Trends Genet 24, 485–488 (2008).

3. Dandage, R. & Landry, C. R. Paralog dependency indirectly affects the robustness of human cells. Mol Syst Biol 15, e8871 (2019).

4. Ohno, S. Evolution by Gene Duplication. (1970) doi:10.1007/978-3-642-86659-3.

5. Aldana, M., Balleza, E., Kauffman, S. & Resendiz, O. Robustness and evolvability in genetic regulatory networks. J Theor Biol 245, 433–448 (2007).

6. Muller, F. L. et al. Passenger deletions generate therapeutic vulnerabilities in cancer. Nature 488, 337–342 (2012).

7. Ehrenhöfer-Wölfer, K. et al. SMARCA2-deficiency confers sensitivity to targeted inhibition of SMARCA4 in esophageal squamous cell carcinoma cell lines. Sci Rep-uk 9, 11661 (2019).

8. Hoffman, G. R. et al. Functional epigenetics approach identifies BRM/SMARCA2 as a critical synthetic lethal target in BRG1-deficient cancers. Proc National Acad Sci 111, 3128–3133 (2014).

9. Oike, T. et al. A Synthetic Lethality–Based Strategy to Treat Cancers Harboring a Genetic Deficiency in the Chromatin Remodeling Factor BRG1. Cancer Res 73, 5508–5518 (2013).

10. Helming, K. C. et al. ARID1B is a specific vulnerability in ARID1A-mutant cancers. Nat Med 20, 251–254 (2014).

11. Lelij, P. van der et al. Synthetic lethality between the cohesin subunits STAG1 and STAG2 in diverse cancer contexts. Elife 6, e26980 (2017).

12. Benedetti, L., Cereda, M., Monteverde, L., Desai, N. & Ciccarelli, F. D. Synthetic lethal interaction between the tumour suppressor STAG2 and its paralog STAG1. Oncotarget 5, 37619–37632 (2014).

13. Kegel, B. D., Quinn, N., Thompson, N. A., Adams, D. J. & Ryan, C. J. Comprehensive prediction of synthetic lethality between paralog pairs in cancer cell lines. BioRxiv (2020) doi:10.1101/2020.12.16.423022.

14. Tsherniak, A. et al. Defining a Cancer Dependency Map. Cell 170, 564–576.e16 (2017).

15. Parrish, P. C. R. et al. Discovery of synthetic lethal and tumor suppressive paralog pairs in the human genome. Biorxiv 2020.12.20.423710 (2020) doi:10.1101/2020.12.20.423710.

16. Gonatopoulos-Pournatzis, T. et al. Genetic interaction mapping and exon-resolution functional genomics with a hybrid Cas9-Cas12a platform. Nat Biotechnol 38, 638–648 (2020).

17. Dede, M., McLaughlin, M., Kim, E. & Hart, T. Multiplex enCas12a screens detect functional buffering among paralogs otherwise masked in monogenic Cas9 knockout screens. Genome Biol 21, 262 (2020).

18. Kegel, B. D. & Ryan, C. J. Paralog buffering contributes to the variable essentiality of genes in cancer cell lines. Plos Genet 15, e1008466 (2019).

19. Thompson, N. A. et al. Combinatorial CRISPR screen identifies fitness effects of gene paralogues. Nat Commun 12, 1302 (2021).

20. Viswanathan, S. R. et al. Genome-scale analysis identifies paralog lethality as a vulnerability of chromosome 1p loss in cancer. Nat Genet 50, 937–943 (2018).

21. Duijf, P. H. G., Schultz, N. & Benezra, R. Cancer cells preferentially lose small chromosomes. Int J Cancer 132, 2316–2326 (2013).

22. Li, C. H. et al. Sex differences in oncogenic mutational processes. Nat Commun 11, 4330 (2020).

23. Wright, D. J. et al. Genetic variants associated with mosaic Y chromosome loss highlight cell cycle genes and overlap with cancer susceptibility. Nat Genet 49, 674–679 (2017).

24. Kaneko, S. & Li, X. X chromosome protects against bladder cancer in females via a KDM6A-dependent epigenetic mechanism. Sci Adv 4, eaar5598 (2018).

25. Spatz, A., Borg, C. & Feunteun, J. X-Chromosome Genetics and Human Cancer. Nat Rev Cancer 4, 617–629 (2004).

26. Hunter, S., Gramlich, T., Abbott, K. & Varma, V. Y chromosome loss in esophageal carcinoma: An in situ hybridization study. Genes Chromosomes Cancer 8, 172–177 (1993).

27. Agahozo, M. C. et al. Loss of Y-Chromosome during Male Breast Carcinogenesis. Cancers 12, 631 (2020).

28. Minner, S. et al. Y chromosome loss is a frequent early event in urothelial bladder cancer. Pathology 42, 356–359 (2010).

29. Lin, S.-H. et al. Mosaic chromosome Y loss is associated with alterations in blood cell counts in UK Biobank men. Sci Rep-uk 10, 3655 (2020).

30. Forsberg, L. A. et al. Mosaic loss of chromosome Y in peripheral blood is associated with shorter survival and higher risk of cancer. Nat Genet 46, 624–628 (2014).

31. Thompson, D. J. et al. Genetic predisposition to mosaic Y chromosome loss in blood. Nature 575, 652–657 (2019).

32. Guo, X. et al. Mosaic loss of human Y chromosome: what, how and why. Hum Genet 139, 421–446 (2020).

33. Lau, Y.-F. C. Y chromosome in health and diseases. Cell Biosci 10, 97 (2020).

34. Shi, J. et al. Discovery of cancer drug targets by CRISPR-Cas9 screening of protein domains. Nat Biotechnol 33, 661–7 (2015).

35. Dass, B. et al. Loss of polyadenylation protein τCstF-64 causes spermatogenic defects and male infertility. Proc National Acad Sci 104, 20374–20379 (2007).

36. Romeo, V., Griesbach, E. & Schümperli, D. CstF64: Cell Cycle Regulation and Functional Role in 31 End Processing of Replication-Dependent Histone mRNAs. Mol Cell Biol 34, 4272–4284 (2014).

37. Yao, C. et al. Overlapping and distinct functions of CstF64 and CstF64τ in mammalian mRNA 31 processing. Rna 19, 1781–1790 (2013).

38. Youngblood, B. A., Grozdanov, P. N. & MacDonald, C. C. CstF-64 supports pluripotency and regulates cell cycle progression in embryonic stem cells through histone 3⍰ end processing. Nucleic Acids Res 42, 8330–8342 (2014).

39. Minvielle-Sebastia, L., Winsor, B., Bonneaud, N. & Lacroute, F. Mutations in the yeast RNA14 and RNA15 genes result in an abnormal mRNA decay rate; sequence analysis reveals an RNA-binding domain in the RNA15 protein. Mol Cell Biol 11, 3075–3087 (1991).

40. Youngblood, B. A. & MacDonald, C. C. CstF-64 is necessary for endoderm differentiation resulting in cardiomyocyte defects. Stem Cell Res 13, 413–421 (2014).

41. Mohr, S. E., Smith, J. A., Shamu, C. E., Neumüller, R. A. & Perrimon, N. RNAi screening comes of age: improved techniques and complementary approaches. Nat Rev Mol Cell Bio 15, 591–600 (2014).

42. Meyers, R. M. et al. Computational correction of copy number effect improves specificity of CRISPR– Cas9 essentiality screens in cancer cells. Nat Genet 49, 1779–1784 (2017).

43. Behan, F. M. et al. Prioritization of cancer therapeutic targets using CRISPR–Cas9 screens. Nature 568, 511–516 (2019).

44. McDonald, E. R. et al. Project DRIVE: A Compendium of Cancer Dependencies and Synthetic Lethal Relationships Uncovered by Large-Scale, Deep RNAi Screening. Cell 170, 577–592.e10 (2017).

45. Nusinow, D. P. et al. Quantitative Proteomics of the Cancer Cell Line Encyclopedia. Cell 180, 387–402.e16 (2020).

46. Ghandi, M. et al. Next-generation characterization of the Cancer Cell Line Encyclopedia. Nature 569, 503–508 (2019).

47. Fortin, J.-P. et al. Multiple-gene targeting and mismatch tolerance can confound analysis of genome-wide pooled CRISPR screens. Genome Biol 20, 21 (2019).

48. Nijhawan, D. et al. Cancer Vulnerabilities Unveiled by Genomic Loss. Cell 150, 842–854 (2012).

49. Paolella, B. R. et al. Copy-number and gene dependency analysis reveals partial copy loss of wild-type SF3B1 as a novel cancer vulnerability. Elife 6, e23268 (2017).

50. Oughtred, R. et al. The BioGRID database: A comprehensive biomedical resource of curated protein, genetic, and chemical interactions. Protein Sci 30, 187–200 (2021).

51. Modos, D. et al. Identification of critical paralog groups with indispensable roles in the regulation of signaling flow. Sci Rep-uk 6, 38588 (2016).

52. Neggers, J. E. et al. Synthetic Lethal Interaction between the ESCRT Paralog Enzymes VPS4A and VPS4B in Cancers Harboring Loss of Chromosome 18q or 16q. Cell Reports 33, 108493 (2020).

53. Szymańska, E. et al. Synthetic lethality between VPS4A and VPS4B triggers an inflammatory response in colorectal cancer. Embo Mol Med 12, e10812 (2020).

54. Wu, J. et al. Cryo-EM Structure of the Human Ribonuclease P Holoenzyme. Cell 175, 1393–1404.e11 (2018).

55. Welting, T. J. M., Kikkert, B. J., Venrooij, W. J. van & Pruijn, G. J. M. Differential association of protein subunits with the human RNase MRP and RNase P complexes. Rna 12, 1373–1382 (2006).

56. Guerrier-Takada, C., Eder, P. S., Gopalan, V. & Altman, S. Purification and characterization of Rpp25, an RNA-binding protein subunit of human ribonuclease P. Rna 8, 290–295 (2002).

57. Goldfarb, K. C. & Cech, T. R. Targeted CRISPR disruption reveals a role for RNase MRP RNA in human preribosomal RNA processing. Gene Dev 31, 59–71 (2017).

58. Jones, P. A. Functions of DNA methylation: islands, start sites, gene bodies and beyond. Nat Rev Genet 13, 484–492 (2012).

59. Tukiainen, T. et al. Landscape of X chromosome inactivation across human tissues. Nature 550, 244–248 (2017).

60. Balaton, B. P., Cotton, A. M. & Brown, C. J. Derivation of consensus inactivation status for X-linked genes from genome-wide studies. Biol Sex Differ 6, 35 (2015).

61. Dunford, A. et al. Tumor-suppressor genes that escape from X-inactivation contribute to cancer sex bias. Nat Genet 49, 10–16 (2016).

62. Wernitznig, A. et al. Abstract 3227: CLIFF, a bioinformatics software tool to explore molecular differences between two sets of cancer cell lines. 3227–3227 (2020) doi:10.1158/1538-7445.am2020-3227.

63. Sekiguchi, T., Iida, H., Fukumura, J. & Nishimoto, T. Human DDX3Y, the Y-encoded isoform of RNA helicase DDX3, rescues a hamster temperature-sensitive ET24 mutant cell line with a DDX3X mutation. Exp Cell Res 300, 213–222 (2004).

64. Wang, T. et al. Identification and characterization of essential genes in the human genome. Science 350, 1096–1101 (2015).

65. Zuo, E. et al. CRISPR/Cas9-mediated targeted chromosome elimination. Genome Biol 18, 224 (2017).

66. Adikusuma, F., Williams, N., Grutzner, F., Hughes, J. & Thomas, P. Targeted Deletion of an Entire Chromosome Using CRISPR/Cas9. Mol Ther 25, 1736–1738 (2017).

67. Fong, P. C. et al. Inhibition of Poly(ADP-Ribose) Polymerase in Tumors from BRCA Mutation Carriers. New Engl J Medicine 361, 123–134 (2009).

68. Bryant, H. E. et al. Specific killing of BRCA2-deficient tumours with inhibitors of poly(ADP-ribose) polymerase. Nature 434, 913–917 (2005).

69. Farmer, H. et al. Targeting the DNA repair defect in BRCA mutant cells as a therapeutic strategy. Nature 434, 917–921 (2005).

70. Godfrey, A. K. et al. Quantitative analysis of Y-Chromosome gene expression across 36 human tissues. Genome Res 30, 860–873 (2020).

71. Venkataramanan, S., Calviello, L., Wilkins, K. & Floor, S. N. DDX3X and DDX3Y are redundant in protein synthesis. Biorxiv 2020.09.30.319376 (2020) doi:10.1101/2020.09.30.319376.

72. Szappanos, D. et al. The RNA helicase DDX3X is an essential mediator of innate antimicrobial immunity. Plos Pathog 14, e1007397 (2018).

73. Michlits, G. et al. Multilayered VBC score predicts sgRNAs that efficiently generate loss-of-function alleles. Nat Methods 1–9 (2020) doi:10.1038/s41592-020-0850-8.

74. Li, W. et al. MAGeCK enables robust identification of essential genes from genome-scale CRISPR/Cas9 knockout screens. Genome Biol 15, 554 (2014).

75. Li, H. et al. The Sequence Alignment/Map format and SAMtools. Bioinformatics 25, 2078–2079 (2009).

76. Hofmann, M. H. et al. BI-3406, a potent and selective SOS1::KRAS interaction inhibitor, is effective in KRAS-driven cancers through combined MEK inhibition. Cancer Discov CD-20-0142 (2020) doi:10.1158/2159-8290.cd-20-0142.

77. Colaprico, A. et al. TCGAbiolinks: an R/Bioconductor package for integrative analysis of TCGA data. Nucleic Acids Res 44, e71–e71 (2016).

78. Li, Y., Ge, D. & Lu, C. The SMART App: an interactive web application for comprehensive DNA methylation analysis and visualization. Epigenet Chromatin 12, 71 (2019).

79. Bengtsson, H., Simpson, K., Bullard, J. & Hansen, K. aroma.affymetrix: A generic framework in R for analyzing small to very large Affymetrix data sets in bounded memory. Report 745, Department of Statistics, University of California, Berkeley, 2008 (2008).

80. Mayrhofer, M., Viklund, B. & Isaksson, A. Rawcopy: Improved copy number analysis with Affymetrix arrays. Sci Rep-uk 6, 36158 (2016).

81. Greenman, C. D. et al. PICNIC: an algorithm to predict absolute allelic copy number variation with microarray cancer data. Biostatistics 11, 164–175 (2010).

82. Hörmann, A. et al. RIOK1 kinase activity is required for cell survival irrespective of MTAP status. Oncotarget 9, 28625–28637 (2018).

83. Hu, Y. et al. An integrative approach to ortholog prediction for disease-focused and other functional studies. Bmc Bioinformatics 12, 357 (2011).

84. Chen, W.-H., Lu, G., Chen, X., Zhao, X.-M. & Bork, P. OGEE v2: an update of the online gene essentiality database with special focus on differentially essential genes in human cancer cell lines. Nucleic Acids Res 45, D940–D944 (2016).

