## Supplementary Figures, Supplementary Table Legends for "Interrogation of cancer gene dependencies reveals novel paralog interactions of autosome and sex chromosome encoded genes"

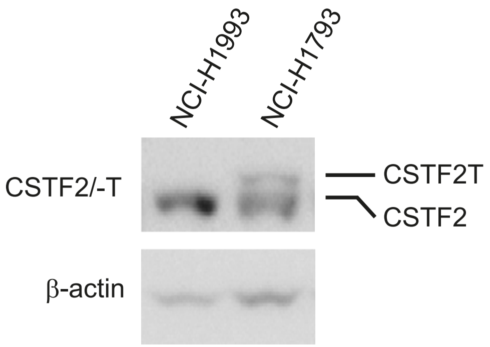


Supplementary Figure 1

Supplementary Figure 1: Related to Figure 1.

Western blot for *CSTF2* and *CSTF2T* in lysates from indicated cell lines. β-actin was used as loading control.

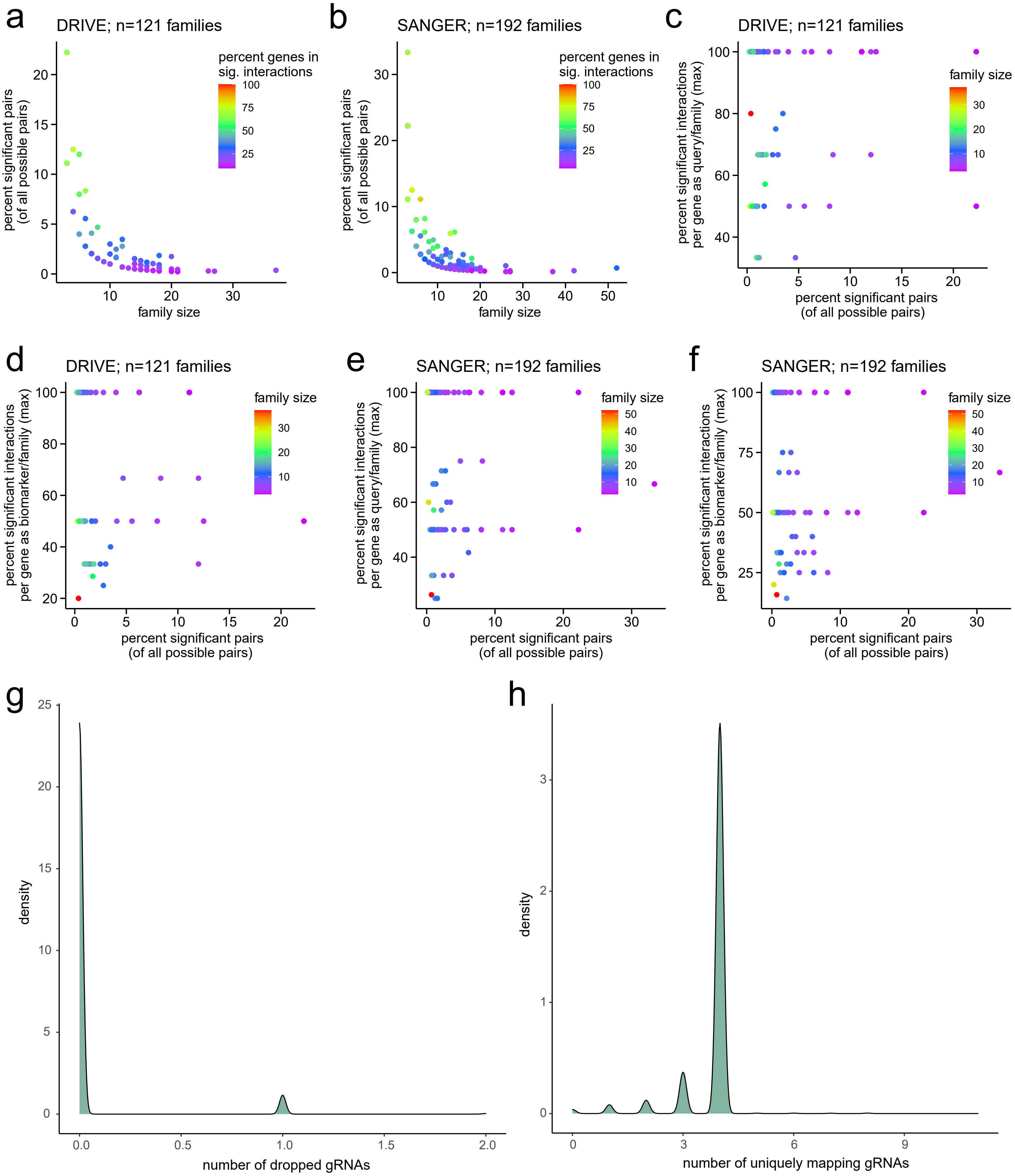
a
DRIVE; n=121 families
percent significant pairs (of all possible pairs)
family size
percent genes in sig. interactions
b
SANGER; n=192 families
c
percent significant interactions per gene as query/family (max)
d
percent significant interactions per gene as biomarker/family (max)
e
f
g
density
number of dropped gRNAs
h
number of uniquely mapping gRNAs

Supplementary Figure 2: Related to Figure 2.

a, b) Percentage of identified hit pairs are plotted against family size from DRIVE (a) and SANGER (b) data. Color indicates percentage of genes from the respective family involved either as query or biomarker.

c-f) Percentage of hit interactions each gene is involved in as query (c, e) or biomarker (d, f) from DRIVE (c, d) and SANGER (e, f) data.

g) Distribution of the number of dropped AVANA gRNAs per query gene.

h) Distribution of the number of uniquely mapped AVANA gRNAs per query gene.

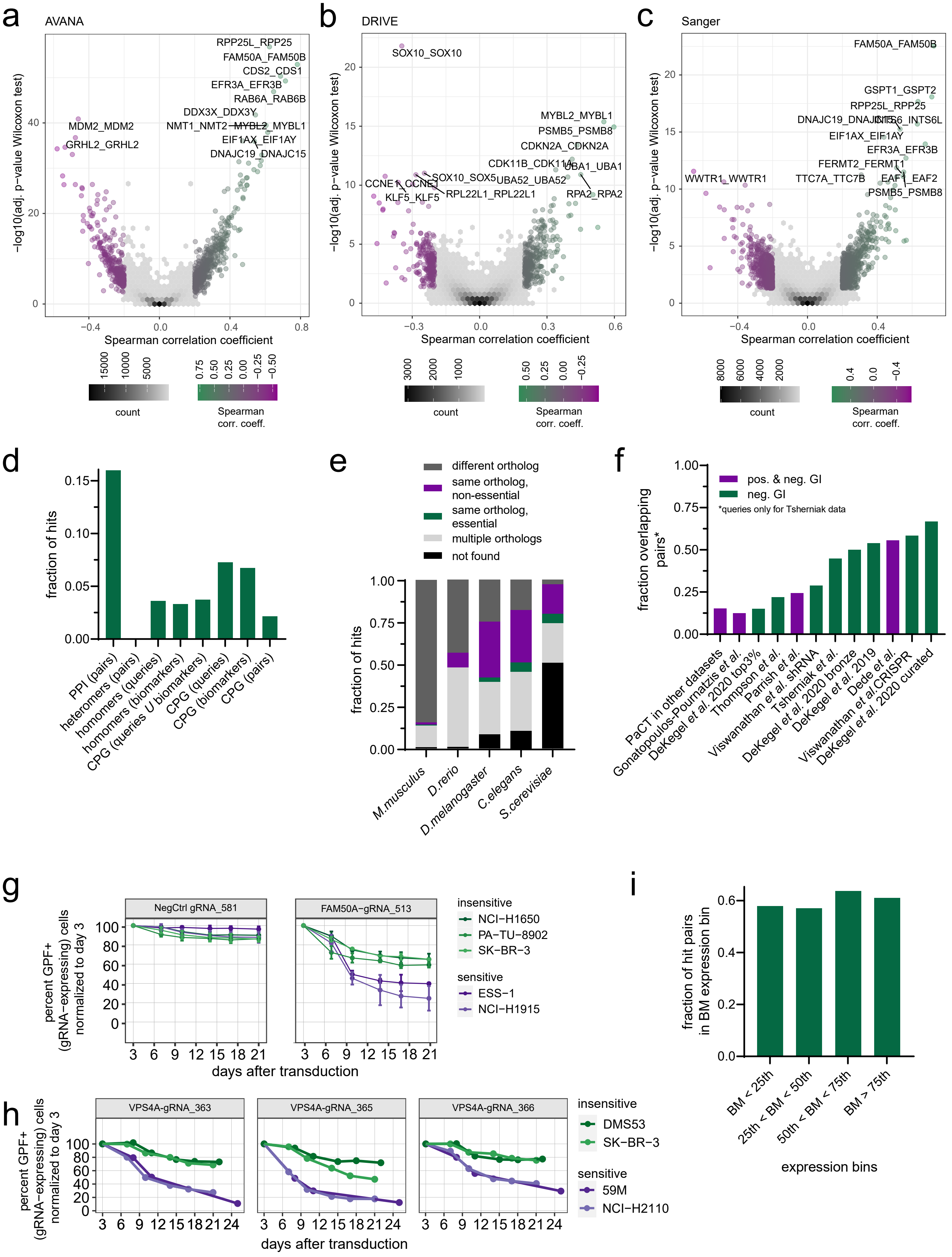


Supplementary Figure 3

Supplementary Figure 3: Related to Figure 3.

a-c) Volcano plots of adjusted Wilcoxon p-value vs. Spearman correlation for AVANA, DRIVE and Sanger data. See Methods for analysis details. Pairs with correlation coefficients <|0.2| and p-values >0.05 are displayed as density plots, 12 most significant pairs are labeled.

d) Annotation of hit pairs: fraction of pairs with known protein-protein-interaction (PPI), with known heteromer interaction or as part of critical paralog groups (CPGs); queries or biomarkers with known homomer interactions or as part of CPGs.

e) Fraction of hit pairs where both paralog genes have the same non-essential ortholog (magenta), the same essential ortholog (green), two different orthologs or multiple orthologs in the indicated model organisms.

f) Fraction of PaCT hit pairs that are recovered in any other dataset (leftmost bar), or of paralog interactions from other datasets identified by PaCT. GI, genetic interactions.

g) CRISPR/Cas9 depletion assay in cell lines resistant (green) and sensitive (purple) to loss of *FAM50A.* gRNAs targeting negative controls (*MP053*) and *FAM50A* are indicated. Cells were lentivirally transduced with the gRNA plasmid containing GFP; GFP percentage in transduced cell line pool was measured by flow cytometry at the indicated time points and normalized to day 3 post-transduction.

h) CRISPR/Cas9 depletion assay in cell lines resistant (green) and sensitive (purple) to loss of *VPS4A.* Additional gRNAs targeting *VPS4A* are indicated (see also Figure 3d). Cells were lentivirally transduced with the gRNA plasmid containing GFP; GFP percentage in transduced cell line pool was measured by flow cytometry at the indicated time points and normalized to day 3 post-transduction.

i) Distribution of PaCT hit pairs across different expression level bins for the biomarker (BM) gene. $25^{th}$, $50^{th}$, $75^{th}$ indicate percentile of expression across all query and biomarker genes in all cell lines included in the PaCT dataset.

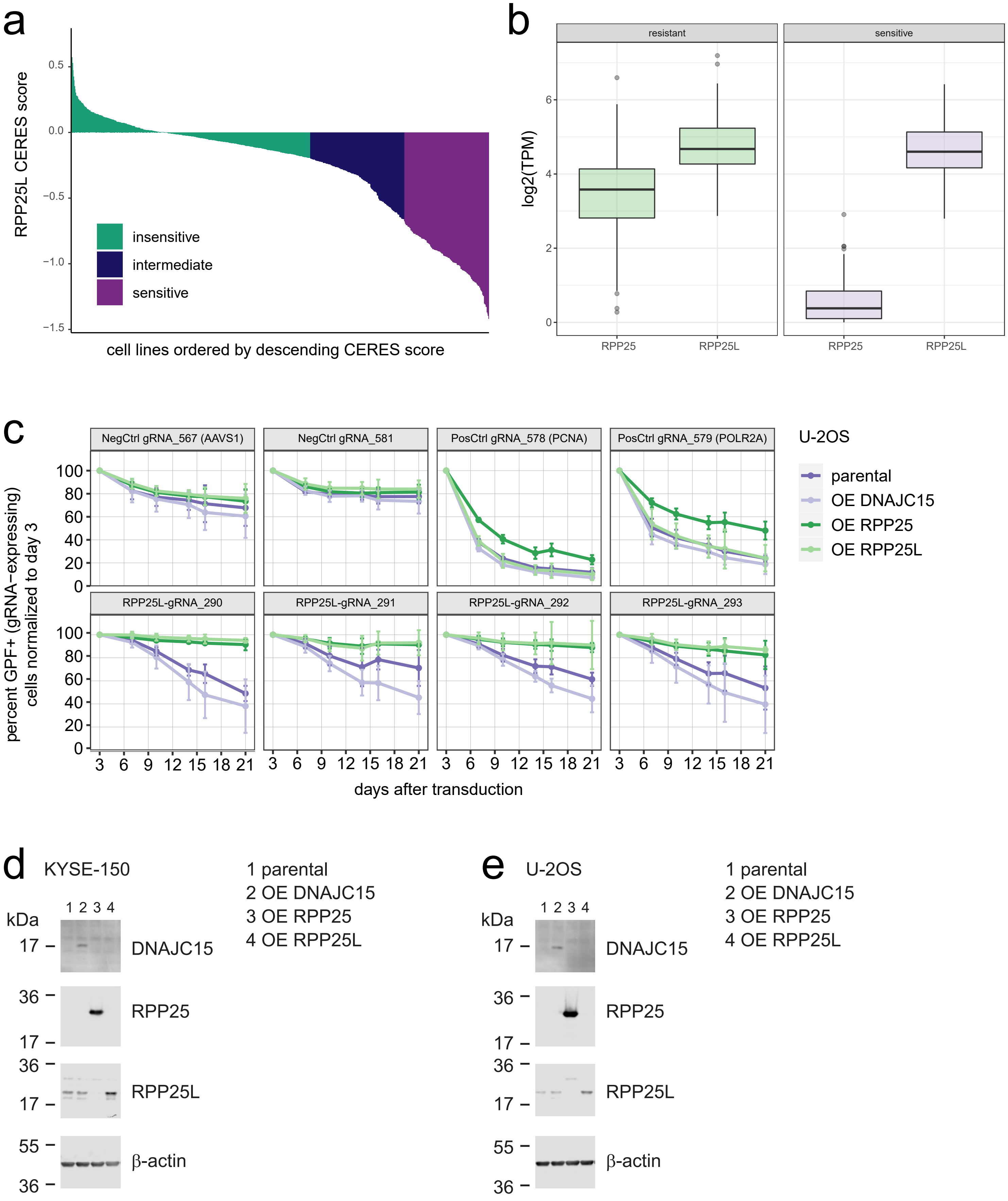
a
RPP25L CERES score
0.5
0.0
−0.5
−1.0
−1.5
insensitive
intermediate
sensitive
cell lines ordered by descending CERES score
b
resistant
sensitive
log2(TPM)
6
4
2
0
RPP25
RPP25L
c
NegCtrl gRNA_567 (AAVS1)
NegCtrl gRNA_581
PosCtrl gRNA_578 (PCNA)
PosCtrl gRNA_579 (POLR2A)
RPP25L-gRNA_290
RPP25L-gRNA_291
RPP25L-gRNA_292
RPP25L-gRNA_293
U-2OS
parental
OE DNAJC15
OE RPP25
OE RPP25L
percent GPF+ (gRNA−expressing) cells normalized to day 3
100
80
60
40
20
0
3 6 9 12 15 18 21
days after transduction
d
KYSE-150
1 parental
2 OE DNAJC15
3 OE RPP25
4 OE RPP25L
kDa
1 2 3 4
17
36
55
DNAJC15
RPP25
RPP25L
β-actin
e
U-2OS
1 parental
2 OE DNAJC15
3 OE RPP25
4 OE RPP25L
kDa
1 2 3 4
DNAJC15
RPP25
RPP25L
β-actin

Supplementary Figure 4: Related to Figure 4.

a) k-means clustering (k=3) separating sensitive, intermediate and resistant cells using AVANA depletion data for *DDX3X*.
b) Boxplot summarizing expression data (log2(TPM)) for members of paralog family in resistant and sensitive cell lines.
c) CRISPR/Cas9 depletion assay in cell lines following ectopic expression of *RPP25* or *RPP25L* in U-2OS cells that are sensitive to loss of *RPP25L* (parental). Overexpression of *DNAJC15* served as a negative control. Expression was induced by addition of doxycycline (0.5 μg/ml) to the medium at the start of the experiment, which was replenished twice per week. gRNAs targeting *RPP25L* (gRNA-290, gRNA-291, gRNA-292, gRNA-293), positive controls (*PCNA*, *POLR2A*) and negative controls (non-targeting, *AAVS1*) are indicated. Cells were lentivirally transduced with the gRNA plasmid containing GFP; GFP percentage in transduced cell line pool was measured by flow cytometry at the indicated time points and normalized to day 3 post-transduction (n=3 independent replicates of the experiment).
d) Western blot for DNAJC15, RPP25 and RPP25L in KYSE-150 cells expressing the indicated overexpression constructs upon culture in the presence of doxycycline (0.5 μg/ml) for 72 hours. $\beta$-actin was included as a loading control.
e) As in (d) for U-2OS cells.

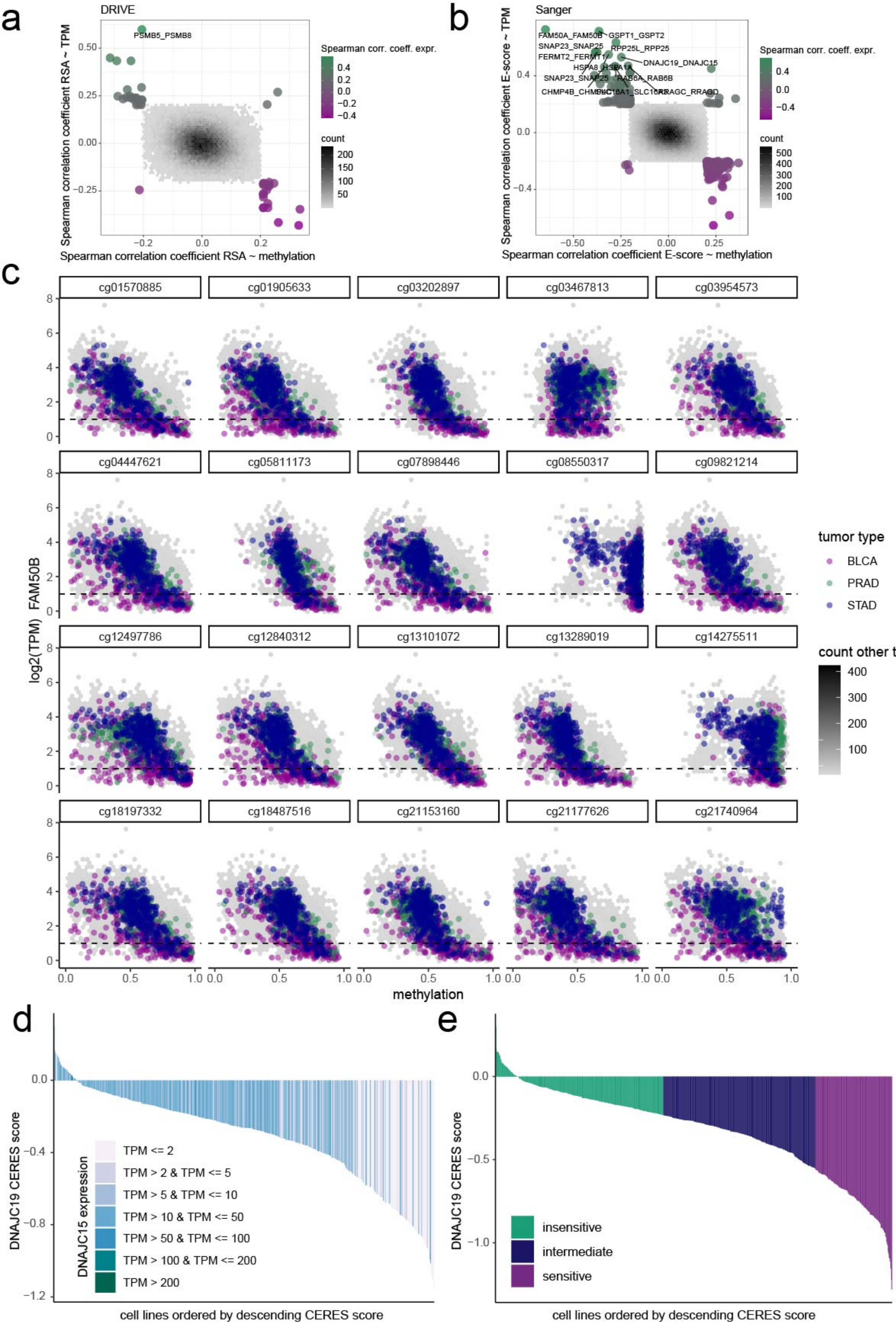


Supplementary Figure 5

Supplementary Figure 5: Related to Figure 5.

a) Scatter plot identifying putative paralog dependencies due to DNA hypermethylation. X-axis: Spearman correlation coefficient between depletion data (RSA score (DRIVE data)) and DNA methylation. Y-axis: Spearman correlation coefficient between depletion data (RSA score (DRIVE data)) and gene expression (TPM). Pairs with correlation coefficients <|0.2| are displayed as density plots, strongest correlations are labeled.
b) As in (a) for Sanger data (E-score).
c) Scatter plot of mRNA expression levels (log2(TPM)) of *FAM50B* versus CpG island methylation across tumour types from TCGA. Samples from bladder urothelial carcinoma (BLCA), prostate adenocarcinoma (PRAD) and stomach adenocarcinoma (STAD) studies are highlighted.
d) AVANA-based depletion scores (CERES) for *DNAJC19,* color-coded by *DDX3Y* expression levels.
e) k-means clustering (k=3) separating sensitive, intermediate and resistant cells using AVANA depletion data for *DNAJC19*.

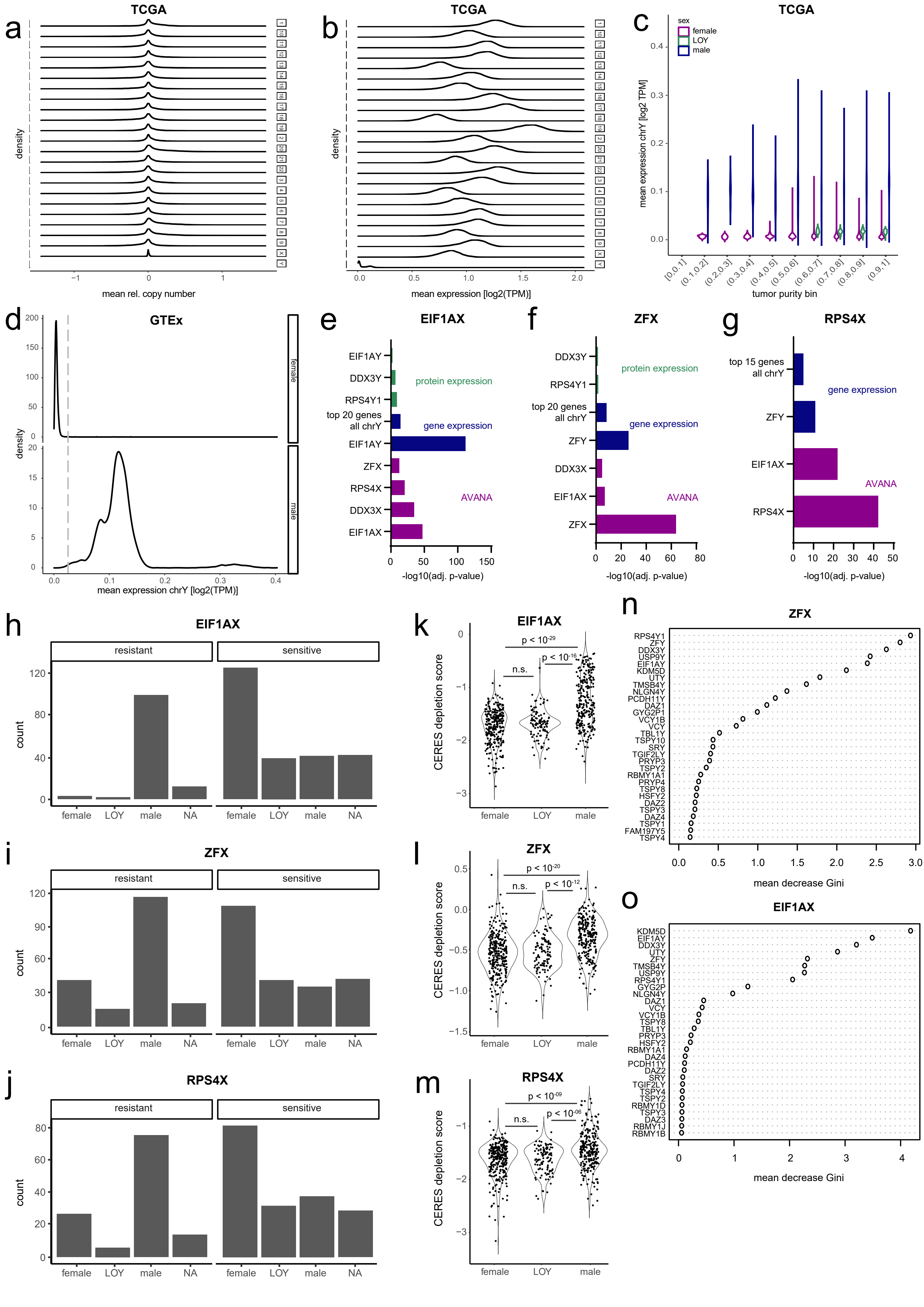


Supplementary Figure 6

Supplementary Figure 6: Related to Figure 6.

a) Distribution of average relative copy number (CN) across genes located on each chromosome for TCGA samples where data were available. CN data for chrY are not available.
b) As in (a) for average gene expression (TPM).
c) Average gene expression (TPM) across genes located on chrY for TCGA samples where data were available. Grouped by sex and by tumour purity bin (as annotated in TCGA). Sex (male, female) as annotated in TCGA or inferred (LOY) as described in Methods.
d) Distribution of average gene expression (TPM) across genes located on chrY for GTEx samples where data were available. Sex (male, female) as annotated in GTEx. No LOY samples were detected by inference.

e-g) Analysis of factors that are most significantly different between *EIF1AX*- (e), *ZFX*- (f), or *RPS4X*- (g) loss-sensitive and resistant cell lines, as defined using k-means clustering based on AVANA data. For each data domain, the most significant discriminators are displayed.

h-j) Sensitive vs. resistant cell lines (as in (e-g)) by sex as annotated in CCLE (male, female) or inferred (LOY) as described in Methods.

k-m) Sensitivity (CERES depletion score for *EIF1AX* (k), *ZFX* (l) or *RPS4X* (m) from AVANA dataset) by sex (as in (h-i)). p-values were calculated using a two-sided Fisher's exact test for count data with Monte-Carlo-simulated p-value (based on 10000 replicates).

n) Variable importance plot for Random Forest model to predict *ZFX* sensitivity. Gene expression values were used as variables for the Indicated genes on y-axis.

o) Variable importance plot for Random Forest model to predict *EIF1AX* sensitivity. Gene expression values were used as variables for the Indicated genes on y-axis.

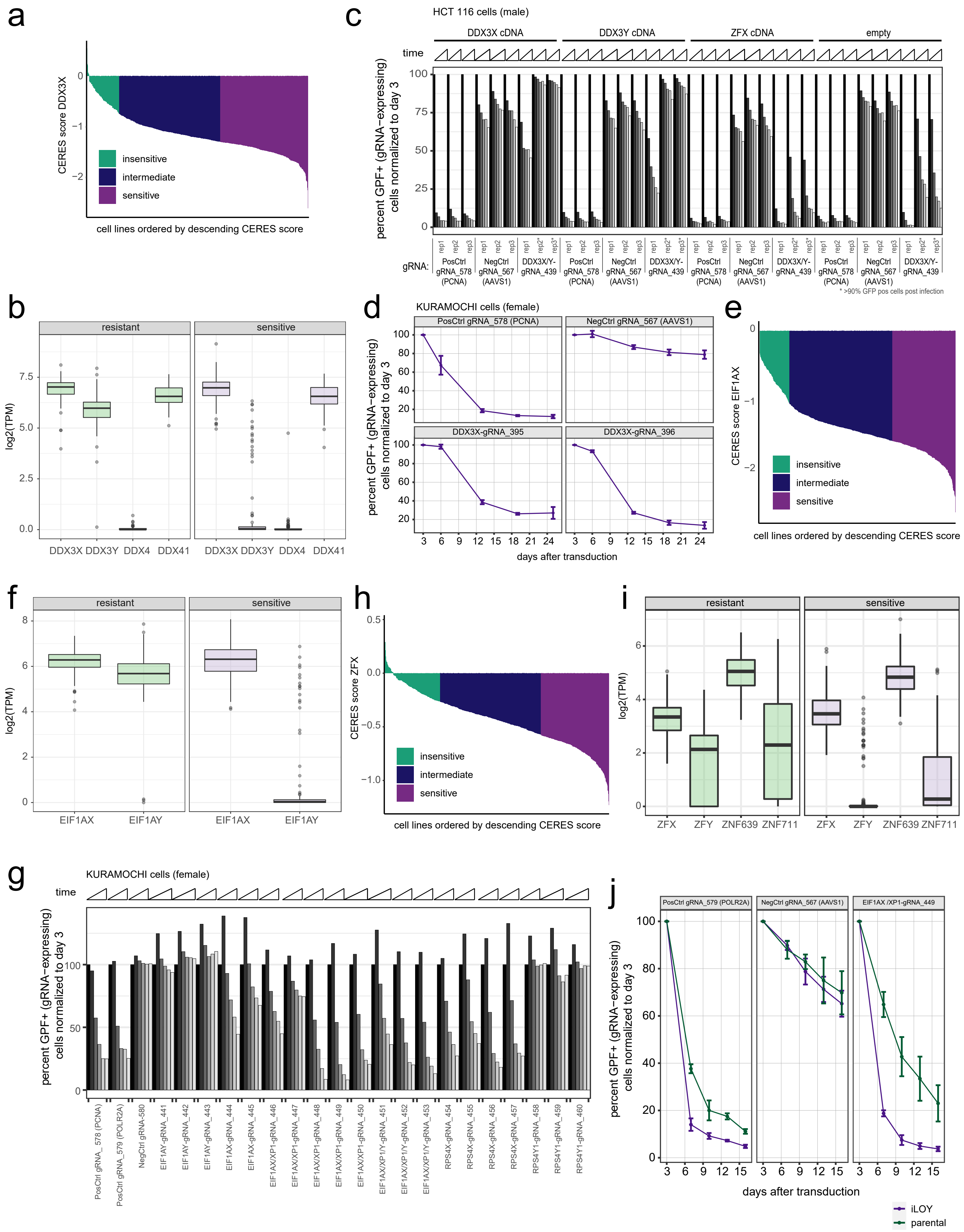


Supplementary Figure 7

Supplementary Figure 7: Related to Figure 7.

a) k-means clustering (k=3) separating sensitive, intermediate and resistant cells using AVANA depletion data for *DDX3X*.
b) Boxplot summarizing expression data ($log_2$(TPM)) for members of *DDX3X* paralog family in resistant and sensitive cells.
c) CRISPR/Cas9 depletion assay in male HCT 116 cells that carry chrY. gRNAs targeting positive control (*PCNA*), negative control (*AAVS1*) and gRNAs targeting *DDX3X* and *DDX3Y* simultaneously are indicated. Cells were lentivirally transduced with the gRNA plasmid containing GFP; GFP percentage in transduced cell line pool was measured by flow cytometry at the indicated time points and normalized to day 3 post-transduction. Cells were additionally transduced with empty vector (control) or unrelated cDNA encoding *ZFX* (control) and rescue constructs with cDNA encoding *DDX3X* or *DDX3Y*. 3 replicates are plotted next to each other.
d) CRISPR/Cas9 depletion assay in KURAMOCHI cells derived from a female patient. gRNAs targeting positive control (*PCNA*), negative control (*AAVS1*) and *DDX3X* are indicated. Assay as in (c), points in line graph represent mean and error bars denote the standard deviation (n= 3 independent experiments).
e) As (a) for *EIF1AX*.
f) As (b) for *EIF1AX*.
g) CRISPR/Cas9 depletion assay in KURAMOCHI cells derived from a female patient transduced with gRNAs targeting positive control (*PCNA, POLR2A*), negative control (*MP-49*) and indicated genes. *EIF1AX/XP1* indicates *EIF1AX* and the *EIF1AXP1* pseudogene. Assay as in (b). Only one replicate but multiple gRNAs targeting the same genes were done in the course of this experiment.
h) As (a) for *ZFX*.
i) As (b) for *ZFX*.
j) CRISPR/Cas9 depletion assay in HT-1080 parental (male) cells and HT-1080 iLOY cells transduced with gRNAs targeting positive control (*POLR2A*), negative control (*AAVS1*) and *EIF1AX* and the *EIF1AXP1* pseudogene. Points in line graph represent mean, and error bars denote the standard deviation (n= 3 independent experiments).

**Supplementary Table Legends**

Supplementary Table 1:

CRISPR library (gRNA sequences); results from proof-of-concept CRISPR paralog deletion screens (escore, FDR); genomic annotation (deletions) of cell lines used.

Supplementary Table 2:

PaCT results for paralog pairs and self-interactions based on gene/protein expression, or correlations with methylation (correlation coefficients, p-values, paralog family ID, paralog family size); number of dropped gRNAs (AVANA) for query genes.

Supplementary Table 3:

chrX/Y STR results (amelogenin marker).

Supplementary Table 4:

siRNA and gRNA details, plasmids, primer sequences, antibodies, cell lines and media used in this study.
